# Ensuring meiotic DNA break formation in the mouse pseudoautosomal region

**DOI:** 10.1101/536136

**Authors:** Laurent Acquaviva, Michiel Boekhout, Mehmet E. Karasu, Kevin Brick, Florencia Pratto, Megan van Overbeek, Liisa Kauppi, R. Daniel Camerini-Otero, Maria Jasin, Scott Keeney

## Abstract

Sex chromosomes in males share only a diminutive homologous segment, the pseudoautosomal region (PAR), wherein meiotic double-strand breaks (DSBs), pairing, and crossing over must occur for correct segregation. How cells ensure PAR recombination is unknown. Here we delineate cis-and trans-acting factors that control PAR ultrastructure and make the PAR the hottest area of DSB formation in the male mouse genome. Prior to DSB formation, PAR chromosome axes elongate, sister chromatids separate, and DSB-promoting factors hyperaccumulate. These phenomena are linked to mo-2 minisatellite arrays and require ANKRD31 protein. We propose that the repetitive PAR sequence confers unique chromatin and higher order structures crucial for DSB formation, X–Y pairing, and recombination. Our findings establish a mechanistic paradigm of mammalian sex chromosome segregation during spermatogenesis.

## Introduction

Meiotic recombination forms connections between homologous chromosomes that ensure accurate segregation (*1*). In many species, every chromosome must recombine, so a crucial challenge is to ensure that every chromosome pair acquires at least one SPO11-generated DSB to initiate recombination (*2*).

This challenge is especially acute in most male placental mammals for sex chromosomes (X and Y), on which a DSB can only support recombination if it occurs in the tiny PAR (*3-7*). The PAR in laboratory mice is the shortest thus far mapped in mammals, at ∼700 kb (*5, 6*). Since only one DSB is formed per ten megabases on average in the mouse, the PAR would risk frequent recombination failure if it behaved like a typical autosomal segment (*7, 8*). However, the PAR is not typical, having disproportionately frequent DSB formation and recombination (*4, 7, 9, 10*). The mechanisms promoting such frequent DSBs are not known in any species.

Higher order chromosome structure plays an important role in meiotic recombination. DSBs arise concomitantly with development of linear axial structures that anchor arrays of chromatin loops within which DSBs occur (*1, 11-13*). The axis begins to form between sister chromatids during pre-meiotic replication (pre-leptonema) and includes SYCP2 and SYCP3 (*14, 15*), cohesin complexes (*16*), and HORMA domain proteins (HORMAD1 and HORMAD2) (*17-20*). Axes are also assembly sites for IHO1, MEI4, and REC114 complexes (*13, 21-24*), whose functions as essential promoters of SPO11 activity are incompletely understood (*25, 26*).

We previously showed that PAR chromatin is organized into short loops on a long axis, (*7*). However, only a low-resolution view of PAR structure was available and the cis-and trans-acting factors controlling PAR structure and DSB formation have remained largely unknown.

## Results

### A distinctive PAR ultrastructure rich in pro-DSB factors

We applied a cytogenetic approach to investigate the mouse PAR structure in the C57BL/6J strain (B6). In most of the genome, axes elongate and DSBs begin to form during leptonema, then homologous chromosomes pair and axes are juxtaposed by the transverse filament protein SYCP1 (forming the synaptonemal complex, or SC) during zygonema. Synapsis and recombination are completed during pachynema, then the SC disassembles during diplonema with homologs remaining attached at sites of crossovers (chiasmata) until anaphase I (*1, 27*). X and Y usually pair late, with PARs paired in less than 20% of spermatocytes at late zygonema when nearly all autosomes are paired (*7, 8*). At this stage, we found by conventional immunofluorescence microscopy that unsynapsed PAR axial elements (SYCP2/3 staining) appeared thickened relative to other unsynapsed axes and had bright HORMAD1/2 staining (**Fig. 1A and fig. S1A,B**) (*28*).

**Fig. 1:**
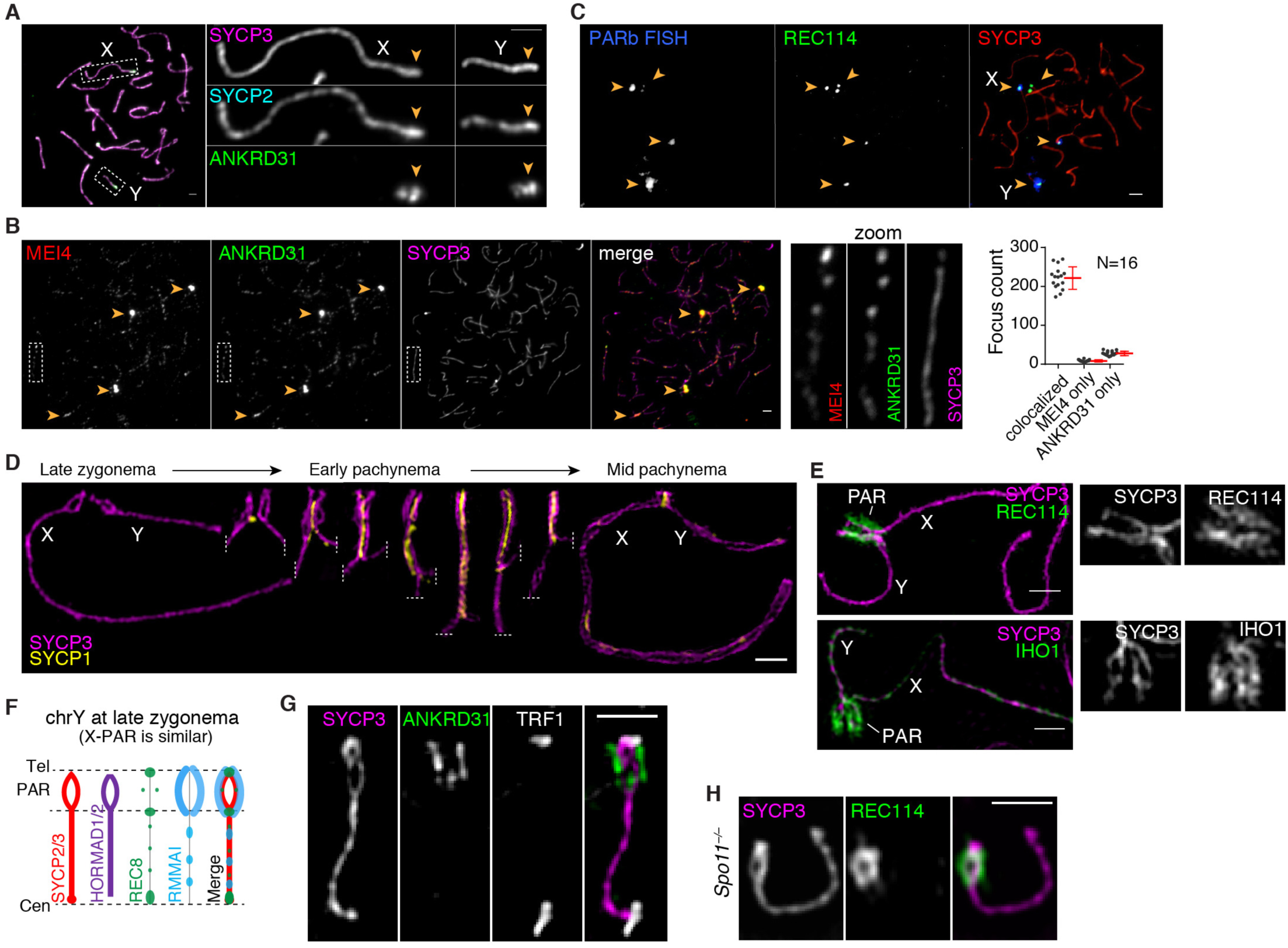
Ultrastructure of the PAR during male meiosis. **(A)** Representative images showing axis thickening (SYCP2 and SYCP3) and ANKRD31 accumulation on X and Y PARs (arrowheads) in late zygonema. Scale bar: 2 μm. **(B)** Colocalization of ANKRD31 and MEI4 in a representative zygotene spermatocyte. Arrowheads indicate blobs. Dashed boxes are shown at higher magnification at the right. Graph: total number of MEI4 and ANKRD31 foci colocalized in leptotene/zygotene spermatocytes. Underlying data for this an all other graphs are provided in **Data File S1**. Scale bar: 2 μm. **(C)** PARb FISH probe colocalizes with REC114 blobs. Scale bar: 2 μm. **(D)** Ultrastructure of the PAR before and after synapsis. A montage of representative SIM images is shown. Dashed lines indicate where chromosomes are cropped. Scale bar: 1 μm. **(E)** RMMAI proteins are enriched along the extended PAR axes. Scale bars: 1 µm. **(F)** Schematic of PAR ultrastructure and distribution of axis and RMMAI proteins at late zygonema. **(G)** Telomere-binding protein TRF1 decorates the tip of the PAR bubble, ruling out a foldback (crozier) configuration. Scale bar: 1 µm. **(H)** PAR axis differentiation occurs in the absence of SPO11-generated DSBs. Scale bar: 1 µm.

We found that the PAR was highly enriched for DSB-promoting factors REC114, MEI4, MEI1, and IHO1 [proteins essential for genome-wide DSB formation (*22-24, 29*)] as well as ANKRD31, a REC114 partner essential for PAR DSB formation (*30, 31*). All five proteins (hereafter RMMAI for simplicity) colocalized in several bright, irregular “blobs” for most of prophase I (**Fig. 1A–C and fig. S1C).** Two blobs were on the X and Y PARs as judged by chromosome morphology at late zygonema (**Fig. 1A**) and particularly bright fluorescence in situ hybridization (FISH) with a probe for the PAR boundary (PARb) (**Fig. 1C**). Other blobs highlighted the distal ends of specific autosomes (**Fig. 1C**), which we revisit below. Undefined blobs are also apparent in published micrographs but were not explored further (*21-24*). Consistent with and extending other studies (*21-24, 30, 31*), all five proteins also colocalized in numerous small foci along unsynapsed chromosome axes (**Fig. 1B and fig. S1C**), but PAR staining was much brighter. ANKRD31, MEI1 and REC114 enrichment on the PAR was already detectable in pre-leptotene cells during premeiotic S phase (**fig. S1D**), as shown for MEI4 and IHO1 (*21, 23*).

Structured illumination microscopy (SIM) resolved the thickened PAR axes as two strands of axial core (**Fig. 1D and fig. S2A,B**) that were heavily decorated along their lengths with RMMAI proteins (**Fig. 1E**). At the zygotene–pachytene transition, X and Y pair and initiate synapsis, then SC spreads bidirectionally—homologously in the PAR and non-homologously between the non-PAR axes, which definitively marks early pachynema in mice (*28, 32, 33*) (**Fig. 1D**). Separation of PAR axes was readily observed by SIM in late zygonema before X and Y pairing and synapsis, remained apparent after synapsis, then disappeared during early pachynema (**Fig. 1D**). A multi-core structure was also seen in earlier electron microscopy studies, but was staged incorrectly as occurring at mid-to-late pachynema (*34*) (**fig. S2C,D**).

In principle, the sister chromatid axes could be split apart (**Fig. 1F**) or the PAR could be folded back on itself in a crozier configuration (**fig. S2E**). However, a crozier was ruled out by using SIM to define the path of the PAR: the telomere binding protein TRF1 (*35*) decorated only the tip of the chromosome (**Fig. 1G**) and the PARb probe yielded FISH signals symmetrically arrayed at similar positions on both axial cores (**fig. S2F**). We conclude that each axial core is one of the sister chromatids, with a “bubble” opened from near the PAR boundary almost to the telomere (**Fig. 1F**). Axis splitting and REC114 enrichment occurred in the absence of SPO11 (**Fig. 1H**), thus neither property of the PAR is provoked by DSB formation.

These findings establish that the X and Y PARs adopt an elaborate axial structure that forms prior to homologous pairing and synapsis and disappears during early pachynema. It normally occurs at or before the time when most cells make PAR DSBs (which is in late zygonema or around the zygotene–pachytene transition (*7*)) but also forms in the absence of DSBs and is preceded by accumulation of high levels of RMMAI proteins, which starts in pre-leptonema.

### Dynamic remodeling of the PAR loop–axis structure

To investigate temporal patterns of axis differentiation, RMMAI composition, and chromatin loop configuration, we used SIM to compare localization of ANKRD31, SYCP3, and the meiotic cohesin subunit REC8 from pre-leptonema to mid-pachynema (**Fig. 2A**). In parallel, we used conventional microscopy and immunoFISH to measure axis lengths (distance from the PARb probe to the end of the SYCP3-labeled axis), REC114 spreading along the axis, and loop sizes defined as axis-orthogonal extension of the PARb FISH signal (**Fig. 2B and fig. S3**). This analysis delineated a dynamic, large-scale reconfiguration of PAR loop–axis structure as prophase I proceeds.

**Fig. 2.**
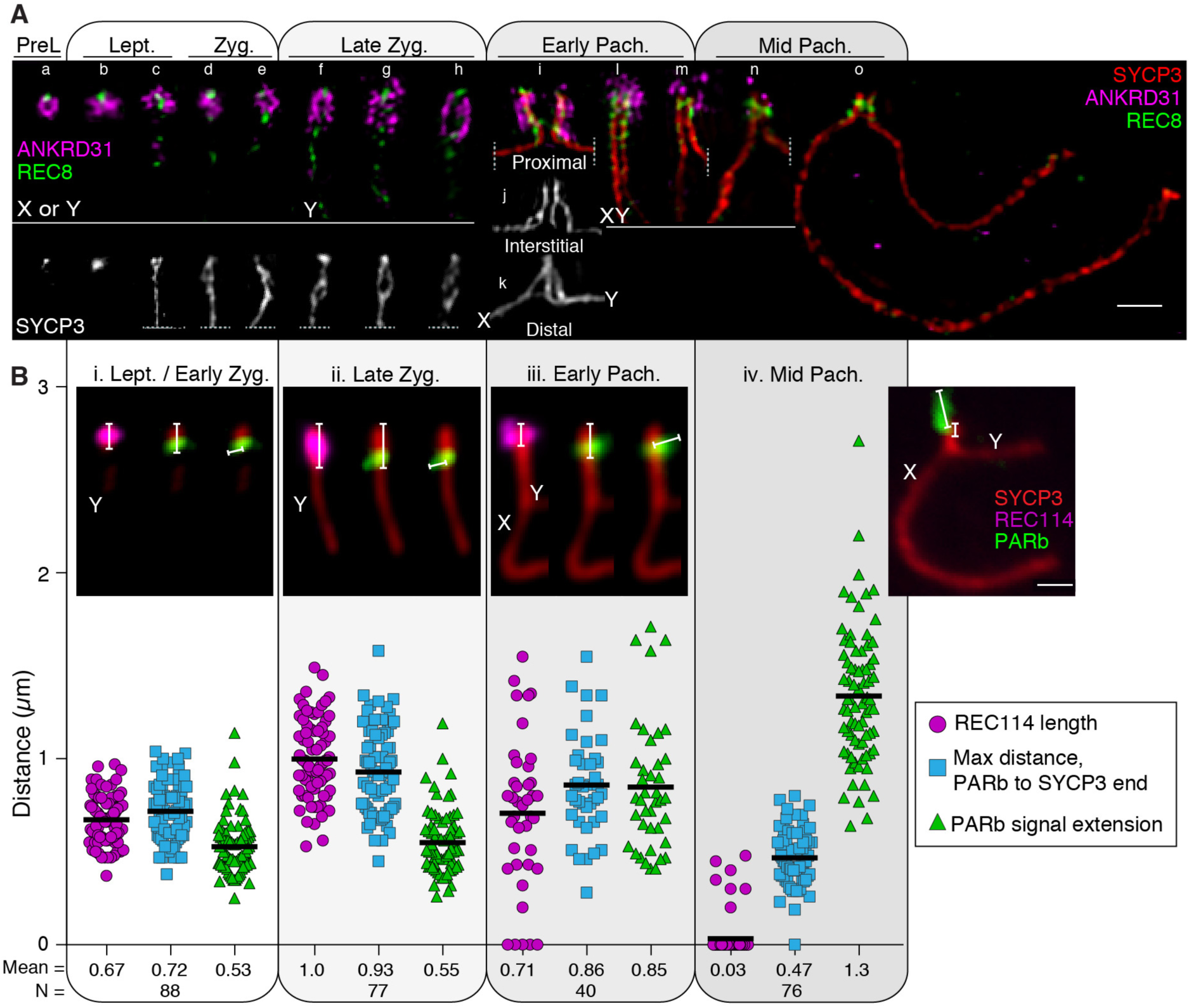
Dynamic remodeling of the PAR loop–axis structure. **(A)** Time course of REC8 and ANKRD31 immunostaining along the PAR axis from pre-leptonema (preL, left) to mid pachynema (right). A montage of representative sex chromosome SIM images is shown. Chromosomes a–e are presumptive X or Y, but could instead be the distal end of chr9. Chromosomes at later stages were unambiguously identified by morphology. Chromosomes i–k show examples where the initial pairing (probably synaptic) contact between X and Y is (i) centromere-proximal (that is, closer to the PAR boundary), (k) distal (that is, closer to the telomere), or (j) interstitial. Scale bar: 1 μm. **(B)** Time course of the spatial organization of the PAR loop–axis ensemble. We collected three measurements from the indicated number (N) of conventional immuno-FISH images from leptonema through mid-pachynema: length of the REC114 signal along the PAR axis; maximal distance from the PARb FISH signal to the distal end of the SYCP3-defined axis; and axis-orthogonal extension of FISH signal for the PARb probe (a proxy for loop sizes). Insets show examples of each type of measurement at each stage. Horizontal black lines on graph indicate means, which are also given numerically below the graph. Quantification from additional mice is provided in **fig. S3**. Scale bar: 1 μm.

At pre-leptonema, ANKRD31 blobs had a closely juxtaposed REC8 focus (**Fig. 2A: chromosome a**). In leptonema and early zygonema, ANKRD31 and REC114 signals stretched along the length of the presumptive PAR axes, while REC8 was restricted to the proximal and distal borders (**Fig. 2A:b–e and 2Bi**). The SYCP3-defined axial element was already long as soon as it was detectable (0.72 µm on average for the mouse in **Fig. 2B**) and the PARb FISH signal was compact (0.53 µm) (**Fig. 2Bi**), confirming and extending our previous observation of long PAR axes and short loops at this stage relative to autosomal loci (*7*).

At late zygonema, the PAR axis had lengthened still further (0.93 µm), while the PARb signal remained compact (**Fig.2Bii**). It was during this stage that the PAR split into separate axial cores, each carrying high levels of RMMAI proteins (**Fig.2A:f–h**). The split corresponded to a REC8-poor zone bounded proximally and distally by focally enriched REC8 (**Fig. 1F and 2A:f–h**).

As cells transitioned into early pachynema and the X and Y PARs synapsed (**Fig. 2A:i–m**), the PAR axes began to shorten slightly (0.86 µm) while the PARb signal expanded (0.85 µm) (**Fig. 2Biii**). Meanwhile, the elongated ANKRD31 signals progressively decreased in intensity, collapsed along with the shortening axes, and separated from the axis while remaining in its vicinity (**Fig. 2A:l–m**). By mid-pachynema, PAR axes collapsed still further, to about half their zygotene length (0.47 µm) and the PARb chromatin expanded to more than twice the zygotene measurement (1.3 µm) (**Fig. 2Biv**). ANKRD31 and REC114 enrichment largely disappeared during this stage, leaving behind a bright bolus of REC8 on the short remaining PAR axis (**Fig. 2A:n–o and 2Biv**).

To sum up, the PAR already has a long axis and compact chromatin loops as soon as the axis forms at the beginning of prophase, and the axis lengthens and sister axes split apart in late zygonema before X and Y pairing. After synapsis, the axes shorten and the chromatin loops decompact. RMMAI proteins are enriched before axis formation and they spread along the axis as it forms and lengthens, then dissociate concomitant with axis collapse in pachynema. This analysis establishes spatial and temporal correlations between presence of RMMAI proteins and the association of compact PARb chromatin with a long axis.

### RMMAI proteins accumulate at mo-2 minisatellites

What are the cis-acting determinants of this PAR behavior? We deduced that DNA sequences shared between the PAR and certain autosomes might be responsible for recruiting RMMAI proteins because the blobs consistently decorated the distal tips of specific autosomes that also hybridized, albeit more weakly, to the PARb FISH probe (**Fig. 1C**).

Self-aligning the PARb probe sequence illustrated its highly repetitive nature, including a central ∼20-kb tandem array of a minisatellite called mo-2, with a 31-bp GC-rich repeat (*36, 37*) (**Fig. 3A**). BLAST searches detected additional mo-2 clusters at the non-centromeric ends of chr9 and chr13 in the mm10 genome assembly and chr4 from the Celera assembly (**Fig. 3A**), matching the reported distribution (*36, 37*). Although these autosomal clusters are only 2 to 8 kb in the assemblies, their copy number is likely underrepresented because they lie adjacent to gaps.

**Fig. 3.**
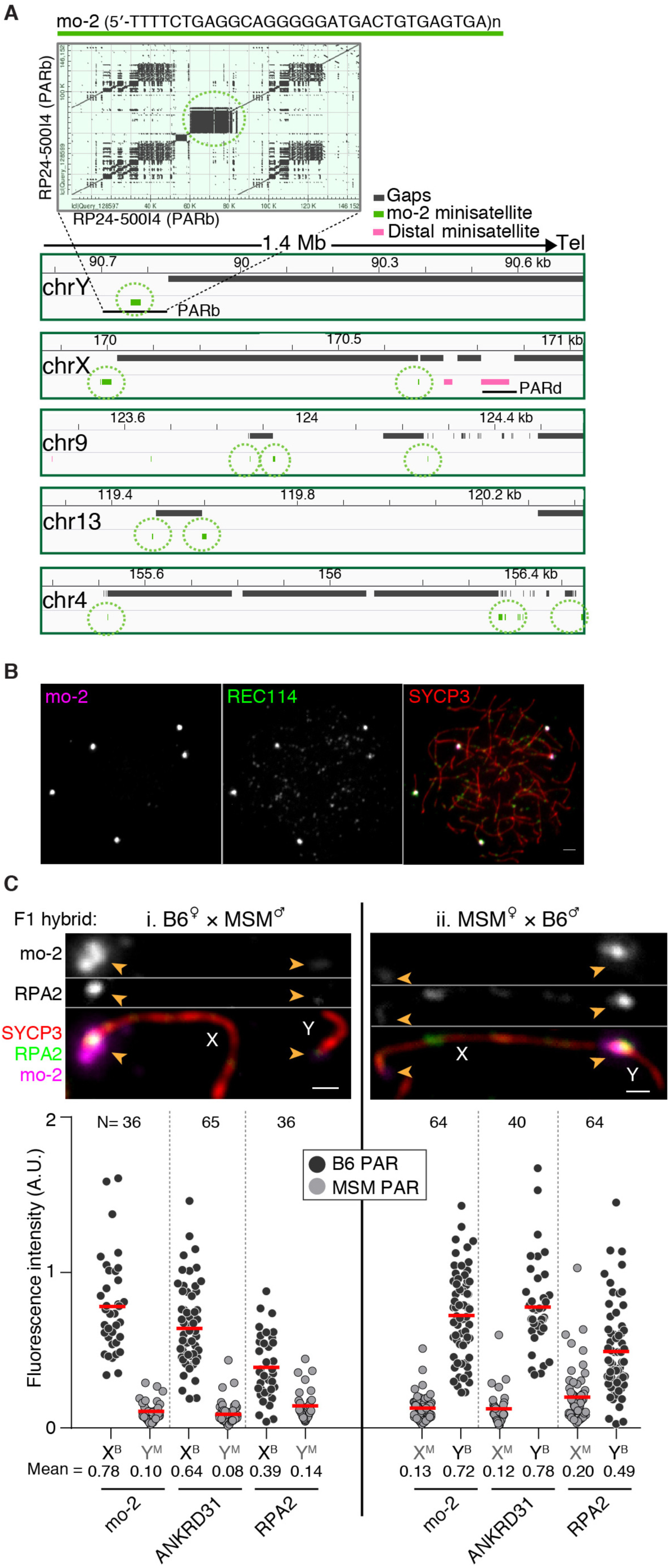
Arrays of the mo-2 minisatellite are sites of RMMAI protein enrichment in the PAR and on autosomes. **(A)** Top panel: Self alignment of the PARb FISH probe, showing its highly repetitive nature. The circled block in the dot plot is a 20-kb cluster of the 31-bp mo-2 minisatellite (*37*). Bottom panel: Schematic depicting the last 1.4 Mb of the non-centromeric ends of the indicated chromosomes, showing the presence of mo-2 repeats (green) adjacent to gaps in chromosome assemblies (mm10). Mo-2 repeats also appear at the distal end of the chr4 in the shotgun assembly from Celera (Mm_Celera, 2009/03/04). The positions of the BAC clones used for FISH are indicated (PARb and PARd). **(B)** Colocalization of REC114 blobs with FISH signal using the mo-2 minisatellite sequence as a probe. A representative zygotene spermatocyte is shown. Scale bar: 2 μm. **(C)** PAR enrichment for ANKRD31 and RPA2 correlates with mo-2 copy number. Top panels: representative micrographs of late zygotene spermatocytes from reciprocal F1 hybrid males from crosses of B6 (high mo-2 copy number) and MSM (low mo-2 copy number) parents. Scale bars: 1 µm. Bottom panels: quantification of PAR-associated signals (A.U., arbitrary units) on B6-derived chromosomes (X^B^ and Y^B^) and MSM-derived chromosomes (X^M^ and Y^M^) from the indicated number of spermatocytes (N). Red lines indicate means. Differences between X and Y PAR intensities are significant for both proteins and for mo-2 FISH in both F1 hybrids (p < 0.0001, paired t-test).

To test if RMMAI blobs correspond to mo-2 arrays, we used an oligonucleotide FISH probe with the mo-2 consensus sequence. This probe gave a compact signal along the PAR axes that was similar to the RMMAI blob pattern in shape and proportional intensity (**Fig. 3B and fig. S4A,B**). We confirmed the identity of the autosomes with chromosome-specific probes (**fig. S4C**).

These findings led us to hypothesize that mo-2 arrays might be a cis-acting determinant of RMMAI recruitment. If so, we reasoned that reducing mo-2 copy number might reduce the amount of RMMAI proteins. To test this prediction, we took advantage of a natural experiment afforded by mouse strain diversity, namely, the fact that the *Mus musculus molossinus* subspecies has substantially lower mo-2 copy number on the PAR and autosomes (*37*). The wild-derived *M. m. molossinus* strain MSM/MsJ (hereafter MSM) showed less hybridization signal with the mo-2 FISH probe as expected, but importantly also gave lower intensity of REC114 immunostaining in blobs compared to B6 (**fig. S4D**).

To quantify this difference and to avoid confounding effects from strain differences in trans-acting factors, we examined spermatocyte spreads from reciprocal F1 hybrid offspring from MSM and B6 parents (**Fig. 3C and fig. S4E**). As we predicted, less ANKRD31 accumulated on MSM-derived PARs. That is, the Y^MSM^ PAR had 8-fold less ANKRD31 than the X^B6^ PAR in F1 hybrids from B6 mothers and MSM fathers (**Fig. 3Ci**), and the X^MSM^ PAR had 6.5-fold less than the Y^B6^ PAR in F1 hybrids from the reciprocal cross (**Fig. 3Cii**). These relative ANKRD31 intensities closely matched relative intensities of mo-2 FISH signals. Despite these quantitative differences, MSM PARs clearly meet the minimal requirements to support sex chromosome pairing with efficiency and timing similar to B6 (**fig. S4F**), not suprisingly since *M. m. molossinus* is a fully fertile subspecies. Interestingly, however, although staining for the ssDNA binding protein RPA was still present on MSM PARs it was nonetheless lower in intensity (**Fig. 3C**). We revisit this observation below.

### Both RMMAI recruitment and PAR loop – axis remodeling require ANKRD31 and MEI4

To identify trans-acting factors important for PAR behavior, we tested mutations eliminating RMMAI components or axis proteins. We counted the number of axial RMMAI foci genome wide and assessed mo-2-associated blobs in leptotene/early-zygotene spermatocytes (**Fig. 4A and fig. S5A**), and examined PAR structure in late zygonema by immuno-FISH for REC114 and mo-2 by conventional microscopy (**Fig. 4B and fig. S5B**) and SIM (**Fig. 4C**).

**Fig. 4.**
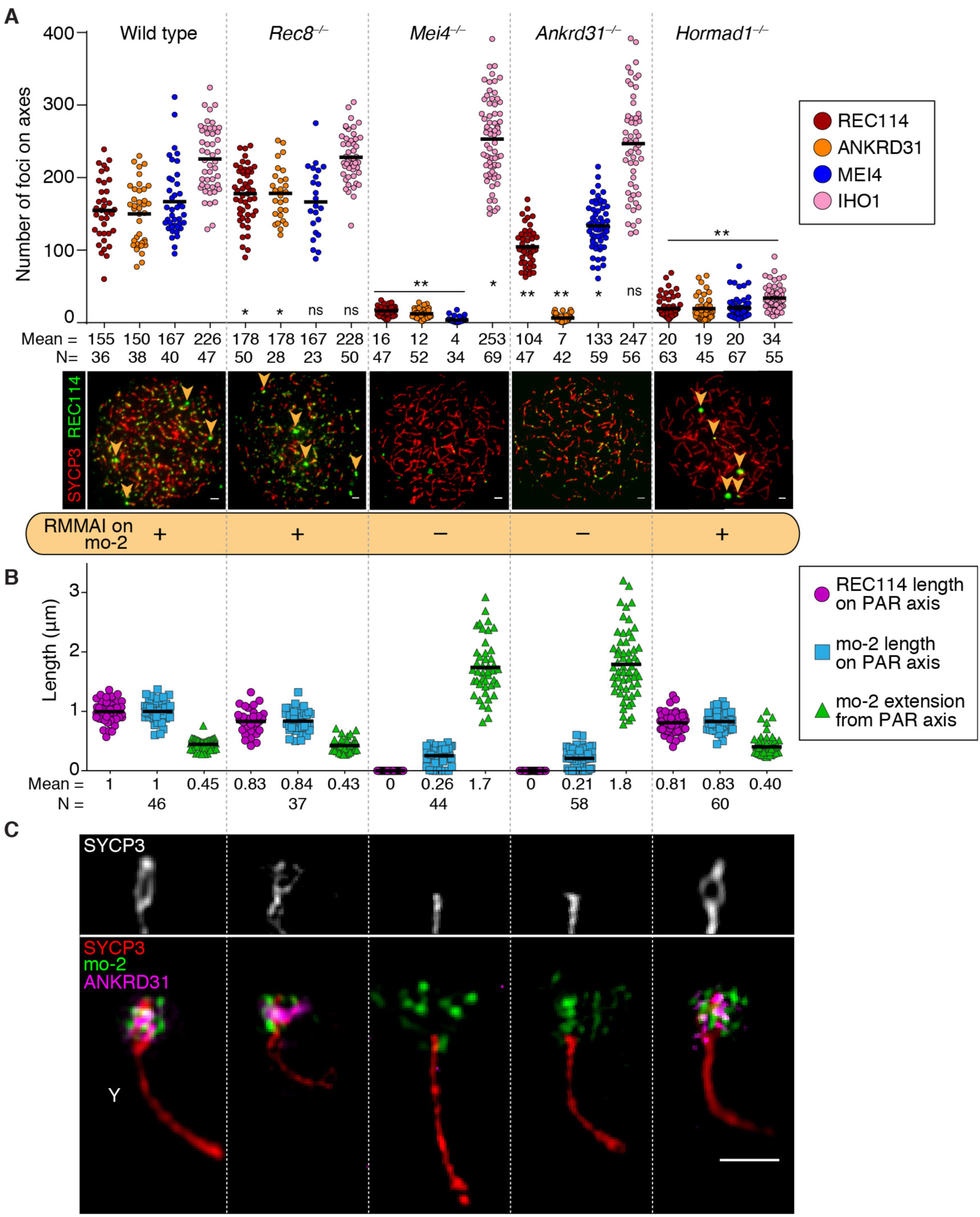
Requirements for RMMAI recruitment and PAR axis remodeling. **(A)** Quantification of REC114, ANKRD31, MEI4, and IHO1 foci along unsynapsed axes in the indicated number of leptotene/early zygotene spermatocytes (N) in wild type and the indicated mutants. Horizontal lines indicate means. Statistical significance for each comparison to wild type is indicated (Student’s t test): * = p<0.02, ** = p≤10^−7^, ns = not significant (p>0.05). Representative micrographs of REC114 staining are shown; images for other proteins are in **fig. S5A**. The presence or absence of mo-2 associated blobs (orange arrowheads) is indicated in the bottom panel for each mutant. Scale bars: 2 μm. **(B)** Genetic requirements for PAR loop–axis organization. Using conventional immuno-FISH microscopy for REC114 and mo-2 in late-zygotene or late-zygotene-like spermatocytes, we measured the length of the REC114 signal along the PAR axis, the length of the mo-2 FISH signal along the PAR axis, and the extension of the mo-2 FISH signal orthogonal to the axis. Quantification from additional mice is provided in **fig. S5B**. **(C)** Representative SIM images showing the Y-PAR loop–axis structure in each mutant at late zygonema. Scale bar: 1 µm.

In *Rec8*^*–/–*^ (*38*), RMMAI foci formed at or slightly above normal numbers along unsynapsed axes [as shown for MEI4 (*21*)] and RMMAI blobs formed on mo-2 regions (**Fig. 4A and fig. S5A**). Moreover, axis elongation, splitting of sister axes, and formation of short loops (i.e., Acquaviva et al., 30 Jan, 2019 – bioRxiv preprint v1 compact mo-2 and REC114 signals) all occurred normally (**Fig. 4B,C and fig. S5B**). However, the PAR axes were separated at the telomeric end and the FISH signals for each sister chromatid in the distal PAR were widely separated (**Fig. 4C and fig. S5C**). We conclude that REC8 maintains cohesion in the distal PAR but is dispensable for RMMAI assembly and formation of the specialized PAR structure.

We tested the importance of RMMAI proteins using *Mei4*^*–/–*^ (*22*) and *Ankrd31*^*–/–*^ (*30*) mutants. In the absence of MEI4, IHO1 foci were not decreased (**Fig. 4A and fig. S5A**). Instead, they were slightly increased, possibly reflecting stabilization if DSBs are not formed (*23*). In contrast, REC114 and ANKRD31 were virtually absent from chromosome axes and did not form blobs on mo-2 regions. In *Ankrd31*^*–/–*^ mice, MEI4 and REC114 foci still formed, but fewer and of lower intensity (**Fig. 4A and fig. S5A,D,E**) (*30*). IHO1 foci formed in high numbers (as in *Mei4*^*–/–*^). Strikingly, however, RMMAI proteins did not accumulate detectably in mo-2-associated blobs (**Fig. 4A and fig. S5A**). Moreover, the normal PAR ultrastructure was absent in both *Mei4*^*–/–*^ and *Ankrd31*^*–/–*^ mutants: axes were short and showed no sign of splitting, and the mo-2 FISH signal was highly extended (**Fig. 4B,C and fig. S5B**). MEI4 is thus essential for assembly of RMMAI foci genome wide, while ANKRD31 contributes genome-wide but is more specifically essential for high-level enrichment at mo-2 regions. These results agree with findings of Tóth and colleagues, who further observed that REC114 but not IHO1 is essential for formation of blobs of the other RMMAI proteins (*31*).

HORMAD1 is important for formation of MEI4 and IHO1 foci genome wide (*21, 23*). In agreement, we found fewer foci of these and other RMMAI proteins in the *Hormad1*^*–/–*^ mutant (*20*) (**Fig. 4A and S5A**). Remarkably, however, HORMAD1 was dispensable for RMMAI blobs (**Fig. 4A and fig. S5A**) and for PAR axial and chromatin structures (**Fig. 4B,C and fig. S5B**).

Several conclusions emerge. First, PAR RMMAI blobs share genetic requirements with autosomal mo-2 blobs for their formation: *Mei4* and *Ankrd31* are essential but *Rec8* and *Hormad1* are dispensable. Second, these requirements are distinct from those for formation of the smaller, more widely dispersed RMMAI foci, i.e., *Hormad1* is important and *Mei4* even more so, but *Ankrd31* contributes only partially. Finally, we establish a functional correlation between the ability to recruit high levels of RMMAI to the PAR and the ability of the PAR to undergo its normal structural differentiation.

### PAR(-like) axis remodeling is tied to mo-2 presence and copy number

If mo-2 arrays are cis-acting determinants of high-level RMMAI recruitment, and if RMMAI assemblies in turn govern PAR structural dynamics, then we would expect that autosomal mo-2 regions should also be able to form PAR-like structures. Indeed, we observed axis splitting at the distal end of chr9 (**Fig. 5A and fig. S6A**), which has the largest autosomal mo-2 array. Moreover, REC114 immunofluorescence and mo-2 FISH signals showed the PAR-like pattern of extended axes and compact chromatin that was fully dependent on *Ankrd31* (**Fig. 5B**). Thus, mo-2 arrays (and/or mo-2-linked elements) are apparently sufficient for both RMMAI recruitment and axis remodeling.

**Fig. 5.**
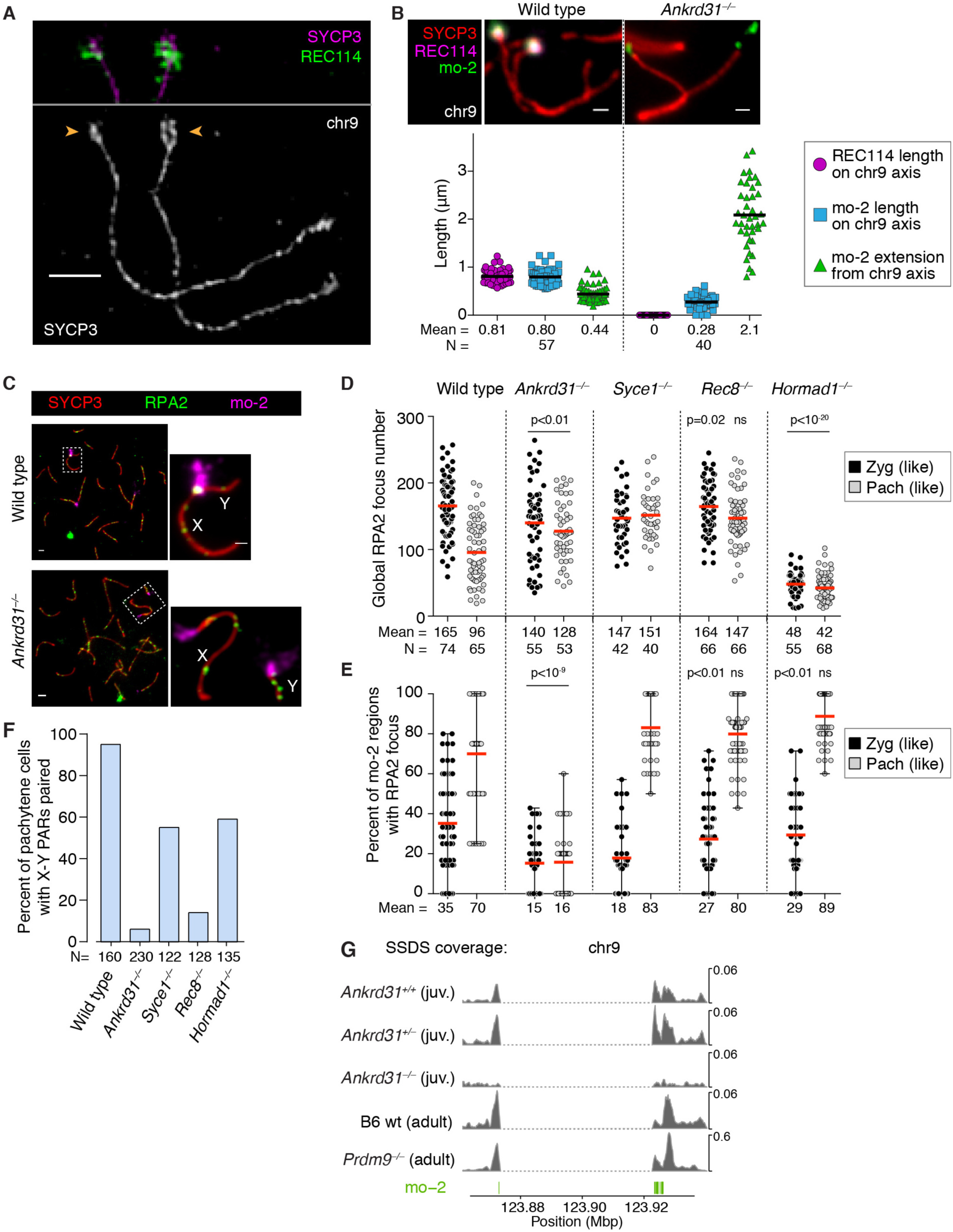
PAR-like structural reorganization and DSB formation on autosomal mo-2 arrays. **(A)** The mo-2-containing portion (arrowheads) of chr9 undergoes axis elongation and splitting similar to PARs. A representative SIM image of a wild-type zygotene spermatocyte is shown. See **fig. S6A** for images of the entire spread and of the Y and PAR-containing part of the X for comparison. Scale bar: 1 µm. **(B)** Loop-axis organization of the mo-2 region of chr9 in wild type and *Ankrd31*^*–/–*^ late-zygotene spermatocytes. Compare to related parameters for the PAR in **Fig. 4B**. Scale bars: 1 µm. **(C–E)** ANKRD31 is required for high-level DSB formation in mo-2 regions in the PAR and on autosomes. Immuno-FISH for RPA2 and mo-2 was used to detect DSBs cytologically in wild type and the indicated mutants. Panel **C** and **fig. S7A** show representative micrographs (scale bar: 2 µm, inset 1 µm). Panel **D** shows global counts of RPA2 foci for zygotene (zyg) or zygotene-like cells and for pachytene (pach) or pachytene-like cells. Red lines are means. Panel **E** quantifies mo-2-associated RPA2 foci in the same cells scored in panel D: for each cell, the fraction of mo-2 regions that had a colocalized RPA2 focus was scored. Red lines are means. Statistical significance is indicated for comparisons (Student’s t tests) of wild type to *Ankrd31*^*–/–*^ or of *Syce1*^*–/–*^ to *Rec8*^*–/–*^ or *Hormad1*^*–/–*^ for matched stages. Note that the number of discretely scorable mo-2 regions in panel E varied from cell to cell depending on pairing status. **(F)** X–Y pairing status, quantified by immuno-FISH for SYCP3 and the PARb probe. **(G)** PAR-like DSB formation near autosomal mo-2 regions. SSDS sequence coverage (data from (*9, 30*)) is shown for the mo-2-adjacent region of chr9. The dashed portion f indicates a gap in the sequence assembly. Positions of mo-2 repeats are shown below. Chr13 is in **fig. S8B** and the PAR is analyzed separately (*30*).

We also examined PARs from *M. m. molossinus*. In keeping with reduced mo-2 and ANKRD31 (**Fig. 3C**), we observed substantially less axis remodeling for MSM-derived PARs in the reciprocal F1 hybrid strains from B6 and MSM parents (**fig. S6B**). These findings reinforce the correlation between mo-2 presence, RMMAI levels, and PAR ultrastructure.

### Determinants of PAR DSB formation in spermatocytes

We hypothesized that RMMAI recruitment (and possibly also the resulting axis remodeling) creates an environment conducive to high-level DSB formation in spermatocytes. This idea predicts that mutations should affect (or not) all of these processes coordinately and that autosomal mo-2 regions should experience PAR-like DSB elevation. These predictions were met.

To evaluate DSB formation, we quantified the number of axial RPA2 foci as a proxy for global DSB numbers, and assessed the proportion of mo-2 FISH signals that contained an RPA2 focus (**Fig. 5C– E and fig. S7A**). Wild-type spermatocytes had a mean of 165 RPA2 foci per cell in zygonema, which declined to 96 foci per cell in pachynema as DSB repair progressed (**Fig. 5D**). In zygonema, RPA2 foci overlapped on average 35% of each cell’s mo-2 regions, increasing to 70% at pachynema (**Fig. 5E**), at which time 95% of spermatocytes had accomplished homologous X–Y pairing as measured by PAR FISH (**Fig. 5F**).

*Ankrd31*^*–/–*^ mutants had a stark reduction in the percentage of mo-2 FISH signals containing an RPA2 focus (**Fig. 5E**) even though global RPA2 focus numbers were only modestly different from wild type at the stages assayed (**Fig. 5D**). [*Ankrd31*^*–/–*^ mutants form substantially fewer RPA2 foci and other cytological DSB markers at leptonema and early zygonema, but not from mid-zygonema on (*30, 31*).] As a consequence, the X and Y chromosomes were paired in only 6% of mid-pachytene spermatocytes (**Fig. 5F**), whereas most *Ankrd31*^*–/–*^ cells pair and synapse all autosomes (*30, 31*).

To analyze mice with *Rec8* and *Hormad1* mutations, we used *Syce1*^*– /–*^ mutants as a point of comparison because they show similar meiotic progression defects without defects in recruitment of RMMAI proteins. SYCE1 is a component of the central element of the SC (*39*). *Rec8* deficiency did not reduce RPA2 focus formation relative to *Syce1*^*–/–*^, either globally on chromosome axes or specifically on mo-2 regions (**Fig. 5D,E**). Despite this, X–Y pairing was substantially reduced in pachytene-like spermatocytes (**Fig. 5F**), likely reflecting that REC8 promotes interhomolog recombination in many species (*40*).

As HORMAD1 is dispensable for RMMAI recruitment to mo-2 regions and for PAR ultrastructure, we predicted that it would also be dispensable for mo-2-associated DSBs. Indeed, *Hormad1*^*–/–*^ spermatocytes had comparable or higher frequencies of RPA2 foci overlapping mo-2 regions (**Fig. 5E and fig. S7B**) and of X–Y pairing as the *Syce1*^*–/–*^ control (**Fig. 5F**). The high frequency of mo-2-associated RPA2 foci was striking given the substantial global reduction in RPA2 foci (**Fig. 5D**) and DSBs (*19*).

Collectively, these findings establish a tight correlation between RMMAI recruitment (which itself correlates with axis remodeling) and high-frequency DSB formation. Further strengthening this correlation, we noted above that MSM-derived PARs display lower RPA2 staining intensity (**Fig. 3C**). Although this result could mean that each DSB in an MSM PAR forms less ssDNA or binds less RPA2 for other reasons, we think it more likely that the lower RPA2 intensity reflects a lesser tendency to make multiple DSBs. Indeed, multiple RPA2 foci could often be resolved by SIM on PARs of B6 spermatocytes (**fig. S7C**), consistent with prior reports of double PAR crossovers (*4*). Furthermore, examples where multiple RPA2 foci were apparent on as-yet unsynapsed Y chromosomes were less frequent in MSM spermatocytes (**fig. S7D**).

To test more directly whether autosomal mo-2 regions experience PAR-like DSB formation, we used maps of ssDNA bound by the strand-exchange protein DMC1 (ssDNA sequencing, or SSDS) (*9, 41, 42*). Separate studies showed that ANKRD31 is critical for high-level DSB formation in the PAR (*30, 31*). We used our SSDS maps from whole-testis samples of juvenile mice (12 days post partum (*30*)) and other SSDS maps to test for PAR-like (i.e., ANKRD31-dependent) DSB signal at autosomal mo-2 regions.

Because of large gaps in the chromosome assemblies, neither the PAR nor the autosomal mo-2 regions can be fully assessed by SSDS or other current deep-sequencing methods. Nonetheless, the region encompassing the mo-2 cluster on chr9 displayed accumulation of SSDS reads that was substantially reduced in the *Ankrd31*^*–/–*^ mutant (**Fig. 5G and fig. S8A**). A modest ANKRD31-dependent peak was also observed near the mo-2 cluster on chr13 (**fig. S8B**). We did not assess chr4 because Sex chromosome recombination available assemblies are too incomplete.

Another hallmark of PAR DSB formation is that it is substantially independent of the histone methyltransferase PRDM9 (*9*). Similarly, SSDS signal around autosomal mo-2 regions was mostly independent of PRDM9 (**Fig. 5G and fig. S8B**). We conclude that autosomal mo-2 regions not only accumulate PAR-like levels of RMMAI proteins and undergo PAR-like axis remodeling in spermatocytes, they frequently form DSBs in a PAR-like manner.

### Behavior of mo-2-containing regions in oocytes

In females, recombination between the two X chromosomes is not restricted to the PAR, so oocytes do not require the same high level of PAR DSB formation as spermatocytes (*43*). We therefore asked whether the PAR undergoes spermatocyte-like structural changes in oocyte meiosis. We detected robust accumulation of RMMAI proteins on mo-2-containing regions in the PARs and autosome ends in oocytes from leptonema to pachynema (**Fig. 6A,B and fig. S9A**), also consistent with conventional micrographs showing blobs of MEI4 and ANKRD31 (*22, 31*). Similar to spermatocytes, X PARs in oocytes displayed an extended axis and compact PARb FISH signal from leptonema to zygonema and a transition to a shorter axis and more extended PARb signal in pachynema, with gradual loss of REC114 signal upon synapsis (**Fig. 6A**). However, we were unable to detect obvious thickening (conventional microscopy) or splitting (SIM) of the PAR axis, and REC8 did not accumulate to high levels (**Fig. 6A,B**). Moreover, like the PAR (*43*), autosomal mo-2 regions showed little enrichment for DSB signal in SSDS data from wild-type ovaries (**fig. S8A and S9B**). These findings suggest that hyper-accumulation of (at least some of) the RMMAI proteins is an intrinsic feature of mo-2-associatd regions but also show that the full suite of PAR(-like) structural changes and DSB formation is dependent on cellular context. Oocytes presumably lack one or more critical protein factors or post-translational modifications.

**Fig. 6.**
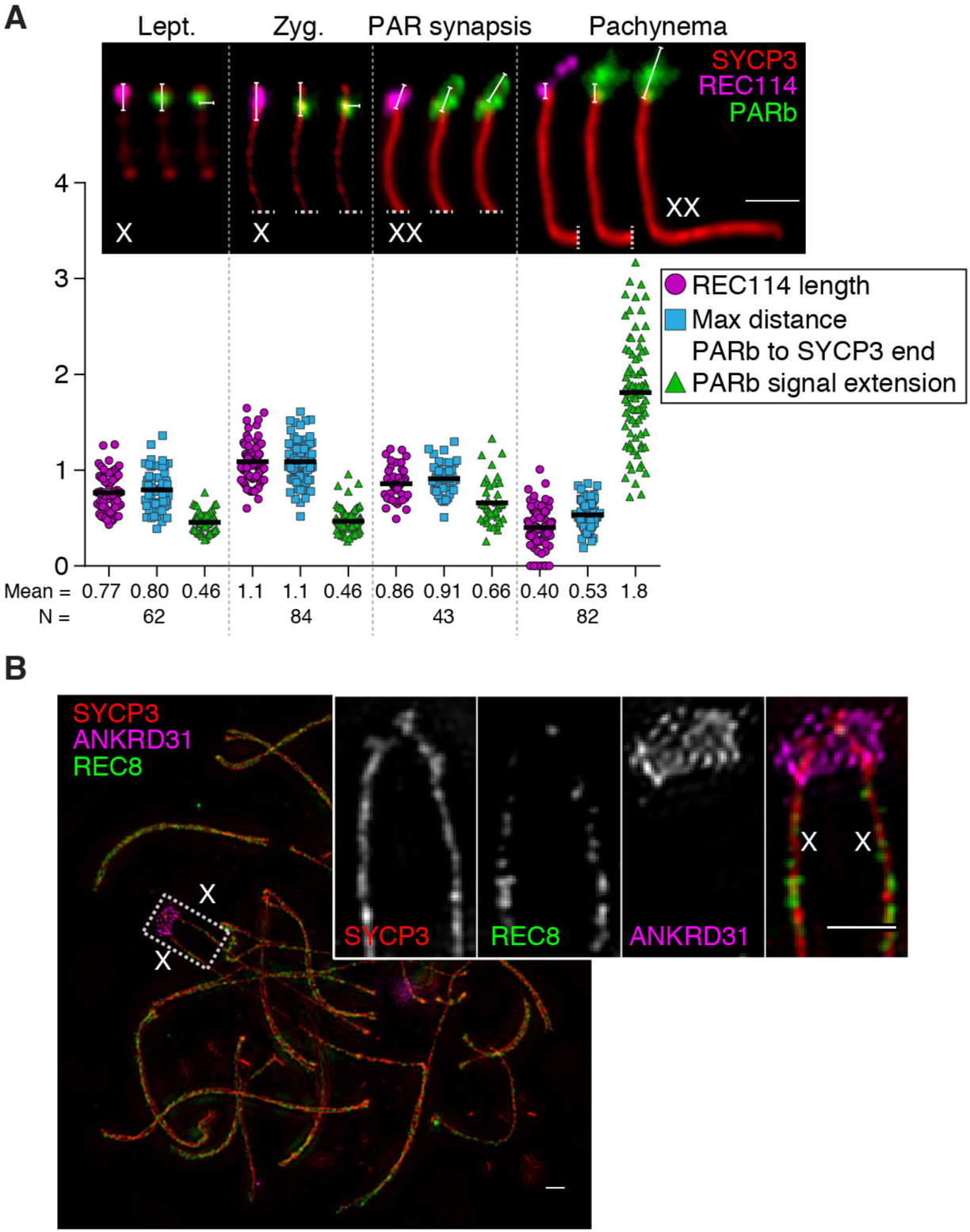
PAR behavior in oocytes. **(A)** PAR ultrastructure in oocytes, quantified as in **Fig. 2B**. Insets show examples of each type of measurement at each stage; late zygotene cells with PAR synapsis are compiled separately from other zygotene cells. Horizontal black lines on graphs indicate means. Scale bar: 1 μm. **(B)** Representative SIM image of a wild-type late zygotene oocyte showing neither detectable splitting of the PAR axis nor REC8 enrichment. Scale bar 2 µm.

## Discussion

We demonstrate that the PAR in male mice undergoes a striking rearrangement of loop–axis structure prior to DSB formation and homologous pairing involving recruitment of RMMAI proteins, dynamic axis elongation, and splitting of sister chromatid axes (f**ig. S10**). Some but not all of these behaviors also occur in oocytes. It appears that the mo-2 minisatellite array is a key cis-acting determinant and at least some of the RMMAI proteins are crucial trans-acting determinants, with ANKRD31 occupying an especially mo-2-specific (as opposed to global) role. Our findings provide mechanistic and ultrastructural explanation for the long axes and short loops in the PAR (*7*). These PAR behaviors appear to be essential for meiotic pairing, recombination, and segregation of heteromorphic sex chromosomes.

RMMAI enrichment could occur through direct DNA binding by one or more RMMAI proteins to the mo-2 DNA sequence (and/or another tightly linked DNA element) or to an mo-2-associated chromatin structure. MEI4, REC114, and ANKRD31 (but not IHO1) are critical for formation of RMMAI blobs and downstream structural changes [this study and (*31*)], but ANKRD31 is less critical than the others for the smaller, more widely distributed RMMAI foci that are thought to be responsible for generating most DSBs genome wide (*21-24*). Ankyrin repeat domains—from which ANKRD31 derives its name—are often involved in protein-protein interactions (*44*), including with chromatin (*45*). The hyperaccumulation of RMMAI may thus reflect the fact that mo-2 repeats provide a highly multivalent moiety that may be recognized principally by ANKRD31 bound to REC114–MEI4 complexes. However, reliance on this repetitive structure likely imposes substantial instability through unequal exchange (*36, 46*). The PAR DNA structure thus paradoxically is necessary for genome stability by supporting sex chromosome segregation but also promotes the known rapid evolution of mammalian PARs (*6*).

The function of PAR axis splitting remains unclear. Sister chromatids develop individualized axes later in prophase (diplonema or diakinesis) in many species (*1*), and expansion microscopy of *Drosophila melanogaster* SCs suggests that sister chromatids have structurally distinct axes during pachynema (*47*). Moreover, splitting of sister chromatid axes occurs in cohesion-defective mutants of mouse (*Rec8*^*–/–*^) and *Sordaria macrospora* (*spo76*) (*48, 49*). There is thus ample precedent for individualization of sister chromatids into separate axes, but to our knowledge, splitting of the PAR in late zygonema represents the earliest stage yet documented for this phenomenon in wild-type meiosis in any species.

Previous studies showed that expressing only one of the two major splicing isoforms of *Spo11* (*Spo11β*) or tagging SPO11 with the DNA binding domain of yeast Gal4 confers a specific defect in PAR DSB formation (*7, 50*). Our findings raise the possibility that the specialized properties of the PAR uniquely sensitize it to otherwise subtle defects in SPO11 activity.

It remains to be determined how the unique loop–axis structure of the PAR emerges from RMMAI accumulation and how this structure is related to DSB formation. Interestingly, budding yeast also uses early, robust recruitment of Rec114 and Mer2 (the IHO1 ortholog) as a means to ensure that its smallest chromosomes incur DSBs in every meiosis (*51*). Thus, even if the cis-acting determinants differ between species, such preferential recruitment appears to be an evolutionarily recurrent strategy for mitigating the risk of recombination failure and missegregation that arises when the length of chromosomal homology is limited.

## Materials and Methods

### Mice

Mice were maintained and sacrificed under U.S.A. regulatory standards and experiments were approved by the Memorial Sloan Kettering Cancer Center (MSKCC) Institutional Animal Care and Use Committee. Animals were fed regular rodent chow with *ad libitum* access to food and water. The *Ankrd31* knockout allele (*Ankrd31*^*em*^*1*^*Sky*^) is a single base insertion mutation (+A) in exon 3; its generation and phenotypic characterization is described elsewhere (*30*). Mice with the *Mei4* knockout allele (*22*) were kindly provided by B. de Massy (IGH, Montpellier, France). All other mice were purchased from the Jackson Laboratory: C57BL/6J (stock #00664), MSM/MsJ (stock #003719), B6N(Cg)-*Syce1*^*tm*^*1*^*b(KOMP)Wtsi*^/2J (stock #026719), B6;129S7-*Hormad1*^*tm*^*1*^*Rajk*^/Mmjax (stock #41469-JAX), B6;129S4-*Rec8*^*mei8*^/JcsMmjax (stock #34762-JAX). Mice were genotyped using Direct Tail lysis buffer (Viagen) following the manufacturer’s instructions.

### Generation of REC8 and REC114 antibodies

To produce antibodies against REC8, a fragment of the mouse *Rec8* gene encoding amino acids 36 to 253 (NCBI Reference Sequence: NP_001347318.1) was cloned into pGEX-4T-2 vector. The resulting fusion of the REC8 fragment fused to glutathione S tranferase (GST) was expressed in *E. coli*, affinity purified on glutathione Sepharose 4B, and cleaved with Precision protease. Antibodies were raised in rabbits by Covance Inc. (Princeton NJ) against the purified recombinant REC8 fragment, and antibodies were affinity purified using GST-REC836-253 that had been immobilized on glutathione sepharose by crosslinking with dimethyl pimelimidate; bound antibodies were eluted with 0.1 M glycine, pH 2.5. Purified antibodies were tested in western blots of testis extracts and specificity was validated by immunostaining of spread meiotic chromosomes from wild type and *Rec8*^*–/–*^ mice.

To produce antibodies against REC114, a fragment of the mouse *Rec114* gene encoding a truncated polypeptide lacking the N-terminal 110 amino acids (NCBI Reference Sequence: NP_082874.1) was cloned into pET-19b expression vector. The resulting hexahistidine-tagged REC114_111-259_ fragment was insoluble when expressed in *E. coli*, so the recombinant protein was solubilized and affinity purified on Ni-NTA resin in the presence of 8 M urea. Eluted protein was dialyzed against 100 mM NaH_2_PO_4_, 10 mM Tris-HCl, 6 M urea, pH 7.3 and used to immunize rabbits (Covance Inc.). Antibodies were affinity purified against purified recombinant His_6_-REC114_111-259_ protein immobilized on cyanogen bromide-activated sepharose and eluted in 0.2 M glycine pH 2.5. The affinity purified antibodies were previously used by Stanzione et al. (*23*), who reported detection of a band of appropriate molecular weight in western blots of testis extracts. However, subsequent analysis showed that this band is also present in extracts of *Rec114*^*–/–*^ testes, and thus is non-specific (C. Brun and B. de Massy, personal communication). Importantly, however, Stanzione et al. also reported detection of immunostaining foci on spread meiotic chromosomes similar to findings reported here and by Boekhout et al. (*30*). This immunostaining signal is absent from chromosome spreads prepared from *Rec114*^*–/–*^ mutant mice (C. Brun and B. de Massy, personal communication). Moreover, this immunostaining signal is indistinguishable from that reported using independently generated and validated anti-REC114 antibodies (*24*). We conclude that our anti-REC114 antibodies are highly specific for the cognate antigen when used for immunostaining of meiotic chromosome spreads.

### Chromosome spreads

Testes were dissected and deposited after removal of the tunica albuginea in 1× PBS pH 7.4. Seminiferous tubules were minced using forceps to form a cell suspension. The cell suspension was filtered through a 70-μm cell strainer into a 15 ml Falcon tube pre-coated with 3% (w/v) BSA, and was centrifuged at 1000 rpm for 5 min. The cell pellet was resuspended in 12 ml of 1× PBS for an additional centrifugation step at 1000 rpm for 5 min and the pellet was resuspended in 1 ml of hypotonic buffer containing 17 mM sodium citrate, 50 mM sucrose, 30 mM Tris-HCl pH 8, 5 mM EDTA pH 8, 0.5 mM dithiothreitol (DTT), 10 μl of 100× Halt protease inhibitor cocktail (Thermo Scientific), and incubated for 8 min. Next, 9 ml of 1× PBS was added and the cell suspension was centrifuged at 1000 rpm for 5 min. The cell pellet was resuspended in 100 mM sucrose pH 8 to obtain a slightly turbid cell suspension, and incubated for 10 min. Superfrost glass slides were divided into two squares using an ImmEdge hydrophobic pen (Vector Labs), then 110 μl of 1% paraformaldehyde (PFA) (freshly dissolved in presence of NaOH at 65°C, 0.15% Triton, pH 9.3, cleared through 0.22 μm filter) and 30 μl of cell suspension was added per square, swirled three times for homogenization, and the slides were placed horizontally in a closed humid chamber for 2 h. The humid chamber was opened for 1 h to allow almost complete drying of the cell suspension. Slides were washed in a Coplin jar 2 × 5 min in 1× PBS on a shaker, and 2 min with 0.4% Photo-Flo 200 solution (Kodak), air dried and stored in aluminum foil at –80°C.

Ovaries were extracted from 14.5–18.5 d post-coitum mice, and collected in 1× PBS pH 7.4. After 15 min incubation in hypotonic buffer, the ovaries were placed on a slide containing 30 μl of 100 mM sucrose pH 8, and dissected with forceps to form a cell suspension. The remaining tissues were removed, 110 μl of 1% paraformaldehyde-0.15% Triton was added, and the slides were gently swirled for homogenization, before incubation in a humid chamber as described above for spermatocyte chromosome spreads.

### Immunostaining

Slides of meiotic chromosome spreads were blocked for 30 min at room temperature horizontally in a humid chamber with an excess of blocking buffer containing 1× PBS, pH 7.4 with 0.05% Tween-20, 7.5% (v/v) donkey serum, 0.5 mM EDTA, pH 8.0, and 0.05% (w/v) sodium azide, and cleared by centrifugation at 13,000 rpm for 15 min. Slides were incubated with primary antibody overnight in a humid chamber at 4°C, or for at least 3 hours at room temperature. Slides were washed 3 × 5 min in 1× PBS, 0.05% Tween-20, then blocked for 10 min, and incubated with secondary antibody for 1–2 hours at 37°C in a humid chamber. Slides were washed 3 × 5 min in the dark on a shaker with 1× PBS, 0.05% Tween-20, rinsed in H_2_O, and mounted before air drying with Vectashield (Vector Labs). Antibody dilutions were centrifuged at 13,000 rpm for at least 5 min before use. Primary antibodies used were rabbit and guinea pig anti-ANKRD31 (*30*) (1:200 dilution), rabbit anti-HORMAD2 (Santa Cruz, sc-82192, 1:50), guinea pig anti-HORMAD2 (1:200) and guinea pig anti-IHO1 (1:200) (gifts from A. Toth (Technical University of Dresden)), goat anti-MEI1 (Santa Cruz, sc-86732, 1:50), rabbit anti-MEI4 (gift from B. de Massy, 1:200), rabbit anti-REC8 (this study, 1:100), rabbit anti-REC114 (this study, 1:200), rabbit anti-RPA2 (Santa Cruz, sc-28709, 1:50), goat anti-SYCP1 (Santa Cruz, sc-20837, 1:50), rabbit anti-SYCP2 (Atlas Antibodies, HPA062401, 1:100), mouse anti-SYCP3 (Santa Cruz, sc-74569, 1:100), goat anti-SYCP3 (Santa Cruz, sc-20845, 1:50), rabbit anti-TRF1 (Alpha Diagnostic, TRF12-S, 1:100). Secondary antibodies used were CF405S anti-guinea pig (Biotium, 20356), CF405S anti-rabbit (Biotium, 20420), CF405S anti-mouse (Biotium, 20080), 488 donkey anti-mouse (Life technologies, A21202), 488 donkey anti-rabbit (Life technologies, A21206), 488 donkey anti-goat (Life technologies, A11055), 488 donkey anti-guinea pig (Life technologies, A11073), 568 donkey anti-mouse (Life technologies, A10037), 568 donkey anti-rabbit (Life technologies, A10042), 568 goat anti-guinea pig (Life technologies, A11075), 594 donkey anti-mouse (Life technologies, A21203), 594 donkey anti-rabbit (Life technologies, A21207), 594 donkey anti-goat (Life technologies, A11058), 647 donkey anti-rabbit (Abcam, ab150067), 647 donkey anti-goat (Abcam, ab150131), all at 1:250 dilution.

### ImmunoFISH and DNA probe preparation

All steps were performed in the dark to prevent loss of fluorescence from prior immunostaining. After the last washing step in the immunostaining protocol, slides were placed horizontally in a humid chamber and the chromosome spreads were re-fixed with an excess of 2% (w/v) paraformaldehyde in 1× PBS (pH 9.3) for 10 min at room temperature. Slides were rinsed once in H_2_O, washed for 4 min in 1× PBS, sequentially dehydrated with 70% (v/v) ethanol for 4 min, 90% ethanol for 4 min, 100% ethanol for 5 min, and air dried vertically for 5-10 min. Next, 15 μl of hybridization mix was applied containing the DNA probe(s) in 70% (v/v) deionized formamide (Amresco), 10% (w/v) dextran sulfate, 2× SSC buffer (saline sodium citrate), 1× Denhardt’s buffer, 10 mM EDTA pH 8 and 10 mM Tris-HCl pH 7.4. Cover glasses (22 x 22 mm) were applied and sealed with rubber cement (Weldwood contact cement), then the slides were denatured on a heat block for 7 min at 80°C, followed by overnight incubation (>14 h) at 37°C. Cover glasses were carefully removed using a razor blade, slides were rinsed in 0.1× SSC buffer, washed in 0.4× SSC, 0.3% NP-40 for 5 min, washed in PBS– 0.05% Tween-20 for 3 min, rinsed in H_2_O, and mounted with Vectashield before air drying.

To generate FISH probes, we used the nick translation kit from Abbott Molecular following the manufacturer’s instructions and using CF dye-conjugated dUTP (Biotium), on BAC DNA from the clones RP24-500I4 (maps to the region of the PAR boundary, PARb probe) CH25-592M6 (maps to the distal PAR, PARd probe), RP23-346H16, RP24-136G21, and CH36-200G6 (centromere-distal ends of chr4, chr9, and chr13, respectively). BAC clones were obtained from the BACPAC Resource Center (CHORI). Labeled DNA (500 ng) was precipitated during 30 min incubation at −20°C after adding 5 μl of mouse Cot-1 DNA (Invitrogen), 0.5 volume of 7.5 M ammonium acetate and 2.5 volumes of cold 100% ethanol. After washing with 70% ethanol and air drying in the dark, the pellet was dissolved in 15 μl of hybridization buffer.

Mo-2 oligonucleotide probes were synthetized by Integrated DNA Technologies, with 6-FAM or TYE(tm) 665 fluorophores added to both 5′ and 3′ ends of the oligonucleotide. The DNA sequence was designed based on the previously defined consensus sequence (*37*), and the probe was used at a final concentration of 10 pmol/μl in hybridization buffer without Cot-1 DNA. The Y-chromosome paint probe was purchased from IDLabs and used at 1:30 dilution in hybridization buffer without Cot-1 DNA.

### EdU incorporation

Seminiferous tubules were incubated in DMEM with 10% FCS and 10 µM EdU at 37°C for 1 h for ***in vitro*** labeling. EdU incorporation was detected using the Click-iT EdU Alexa Fluor 647 imaging kit (Invitrogen) according to the manufacturer’s instructions.

### Image acquisition

Images of spread spermatocytes were acquired on a Zeiss Axio Observer Z1 Marianas Workstation, equipped with an ORCA-Flash 4.0 camera and DAPI, CFP, FITC, TEXAS red and Cy5 filter sets, illuminated by an X-Cite 120 PC-Q light source, with either 63×/1.4 NA oil immersion objective or 100×/1.4 NA oil immersion objective. Marianas Slidebook (Intelligent Imaging Innovations) software was used for acquisition.

Structured illumination microscopy (3D-SIM) was performed at the Bio-Imaging Resource Center in Rockefeller University using an **OMX Blaze 3D-SIM super-resolution microscope (Applied Precision)**, equipped with 405 nm, 488 nm and 568 nm lasers, and 100×/1.40 NA UPLSAPO oil objective (Olympus). Image stacks of several μm thickness were taken with 0.125 μm z-steps, and were reconstructed in Deltavision softWoRx 6.1.1 software with a Wiener filter of 0.002 using wavelength specific experimentally determined OTF functions. Slides were prepared and stained as described above, except that chromosomes were spread only on the central portion of the slides, and the slides mounted using 18 × 18 mm coverslips (Zeiss).

### Image analysis

3D-SIM images are shown either as a z-stack using the sum slices function in Fiji/ImageJ, or as a unique slice. The X and/or Y chromosomes were cropped, rotated and further cropped for best display. For montage display, the X and Y chromosomes images were positioned on a black background using Adobe Illustrator. In the instances where the axes of the X and Y chromosomes were cropped, the area of cropping was labeled with a light gray dotted line. Loop/axis measurements, foci counts, and fluorescence intensity quantification were only performed on images from conventional microscopy using the original, unmodified data.

To measure the colocalization between RMMAI proteins, we costained for SYCP3 and ANKRD31 along with either MEI4, REC114, or IHO1, and manually counted the number of ANKRD31 foci overlapping with SYCP3 and colocalizing or not with MEI4, REC114 or IHO1. These counts were performed in 16 spermatocytes from leptonema to early/mid zygonema.

To quantify the total number of RPA2, MEI4, REC114, ANKRD31, and IHO1 foci, single cells were manually cropped and analyzed with semi-automated scripts in Fiji (*52*) as described in detail elsewhere (*30*). Briefly, images were auto-thresholded on SYCP3 staining, which was used as a mask to use ‘Find Maxima’ to determine the number of foci. Images were manually inspected to determine that there were no obvious defects in determining SYCP3 axes, that no axes from neighboring cells were counted, that artifacts were present, or that foci were missed by the script.

To test for colocalization between RPA2 and mo-2 FISH signals, we manually scored the percentage of mo-2 FISH signals colocalizing at least partly with RPA2. Depending on the progression of synapsis during prophase I, between eight and four discrete mo-2 FISH signals could be detected, corresponding to (with increasing signal intensity) the chr4, chr13, chr9, and the PAR (two signals for each when unpaired, or a single signal for each after homologous pairing/synapsis). Notably, the RPA2 focus was most often found in a slightly more centromere-proximal position compared to the bulk of mo-2 FISH signals, and therefore colocalized partly with mo-2 FISH signals. In the case of the PAR, this position corresponds closely to the region of the PAR boundary (PARb probe). Similar trend was observed on autosomal mo-2 clusters.

For estimates of chromatin extension, we measured the maximal axis-orthogonal distance between the FISH signal and the center of the PAR axis, or the centromere-distal axis for chr9 stained by SYCP3. In mutant mice defective for RMMAI protein recruitment in the mo-2 regions, the PAR axis was defined as the nearest SYCP3 segment adjacent to the telomeric SYCP3 signal.

For quantification of RPA2, ANKRD31, REC8, and mo-2 signal intensity in B6 × MSM and MSM × B6 F1 hybrids, late zygotene spermatocytes with at least one RPA2 focus on X or Y PAR were analyzed. We used the elliptic selection tool in Fiji to define a region of interest around the largest signal in the PAR, and the same selection tool was then positioned on the other PAR axis for comparison. The fluorescence intensity was measured as the integrated density with background substraction.

### Prophase I sub-staging and identification of the PAR

Nuclei were staged according to the dynamic behavior of the autosome and sex chromosome axes during prophase I, using SYCP3 staining. Leptonema was defined as having short stretches of SYCP3 but no evidence of synapsis, early/mid-zygonema as having longer stretches of SYCP3 staining and some synapsis, and late zygonema as having fully assembled chromosome axes and substantial (>70%) synapsis. The X and Y chromosomes generally can be identified at this stage, and the PAR axis is distinguishable because it appears thicker than the centromeric end, particularly near the end of zygonema when autosomes are almost fully synapsed. Early pachynema was defined as complete autosomal synapsis, whereas the X and Y chromosomes could display various configuration: i) unsynapsed, with thickened PAR axes, ii) engaged in PAR synapsis, iii) synapsed in the PAR and non-homologously synapsed along the full (or nearly full) Y chromosome axis. Mid pachynema was defined as showing bright signal from autosome axes, desynapsing X and Y axes remaining synapsed only in the PAR, with short PAR axis. During this stage, the autosomes and the non-PAR X and Y axes are initially short and thick, and progressively become longer and thinner. Late pachynema was defined as brighter autosome axes with a characteristic thickening of all autosome ends. The X and Y non-PAR axes are then long and thin and show excrescence of axial elements. Diplonema was defined as brighter axes and desynapsing autosome, associated with prominent thickening of the autosome ends, particularly the centromeric ends. In early diplonema, the non-PAR axes of X and Y chromosomes are still long and thin and progressively condense to form bright axes, associated with bulges. Most experiments were conducted using SYCP3 in combination with a RMMAI protein, which allows easier distinction between synapsing and desynapsing X and Y chromosomes.

By using only SYCP3 staining, the PARs can only be identified unambiguously from the late zygonema-to-early pachynema transition through to diplonema. From pre-leptonema to mid/late-zygonema, the PARs were identified as the two brightest RMMAI signals, the two brightest mo-2 FISH signals, the two brightest PARb FISH signals, or the two FISH signals from the PARd probe. The Y PAR could be distinguished from the X PAR using the PARb probe, as this probe also weakly stains the chromatin of the non-PAR portion of the Y chromosome.

PAR loop/axis measurements in oocytes were performed on two 14.5–15.5 dpc (days post-coitum) (enriched for leptotene and zygotene oocytes) and two 18.5 dpc female fetuses (enriched for pachytene oocytes).

We found significant variability in the X or Y PAR axis length between different animals in our mouse colony maintained in a C57BL/6J congenic background, and even between different C57BL/6J males obtained directly from the Jackson Laboratory. This is in agreement with previous reports about the hypervariable nature of the mo-2 minisatellite and its involvement in unequal crossing over in the mouse (*4, 37, 46, 53, 54*) (mo-2 was also named DXYmov15 or Mov15 flanking sequences). However, the RMMAI signal intensity/elongation and the PAR axis length were always correlated with mo-2 FISH signal intensity. Importantly, despite this variability, mo-2 and RMMAI proteins were enriched in the PAR and autosome ends of all mice analyzed.

### Analysis of SSDS data

SSDS sequencing data were from previously described studies (*9, 30, 43*) and are all available at the Gene Expression Omnibus (GEO) repository under accession numbers GSE35498, GSE99921, GSE118913. To define enrichment values presented in **fig. S8A**, the SSDS coverage was summed across the indicated coordinates adjacent to the mo-2 repeats. A chromosomal mean and standard deviation for chr9 was estimated by dividing the chromosome into 4-kb bins, summing the SSDS coverage in each bin, and calculating the mean and standard deviation after excluding those bins that overlapped a DSB hotspot. The enrichment score was then defined as the difference between the coverage in the mo-2-adjacent region and the chr9 mean coverage, divided by the chr9 standard deviation.

### Statistical analysis

All statistical tests were performed in R (version 3.4.4) (*55*) and RStudio (Version 1.1.442). Negative binomial regression was calculated using the glm.nb function from the MASS package (version 7.3-49) (*56*).

### Data and software availability

Image analysis scripts are available on Github: https://github.com/Boekhout/ImageJScripts.

## End Matter

### Author Contributions and Notes

LA designed and conducted all of the cytogenetic experiments presented and analyzed the data. MEK generated *Ankrd31* mutant mice and anti-ANKRD31 antibodies. MB and MEK provided *Ankrd31* mutant mice and unpublished data. KB and FP performed SSDS and analyzed the data under the supervision of RDC with input from LA and SK. MvO generated REC8 and REC114 antibodies. LK performed initial characterization and provided unpublished data on PAR ultrastructure and cohesin enrichment. MJ and SK designed and supervised the research, analyzed data, and secured funding. LA and SK wrote the manuscript with input from MJ. All authors edited the manuscript. The authors declare no conflict of interest.

## Acknowledgments

We thank Attila Tóth (Technische Universität Dresden, Germany) and Bernard de Massy (Institut Génétique Humaine, Montpellier, France) for antibodies, mice, discussions, and sharing of unpublished information. We thank Alison North and the Bio-Imaging Resource Center at Rockefeller University for assistance with SIM. This work utilized the computational resources of the NIH HPC Biowulf cluster (http://hpc.nih.gov).

## Supplementary Materials

**Other Supplementary Materials for this manuscript include the following:**

Data S1: Excel spreadsheet containing the underlying data for graphs in the figures.

**Fig. S1.**
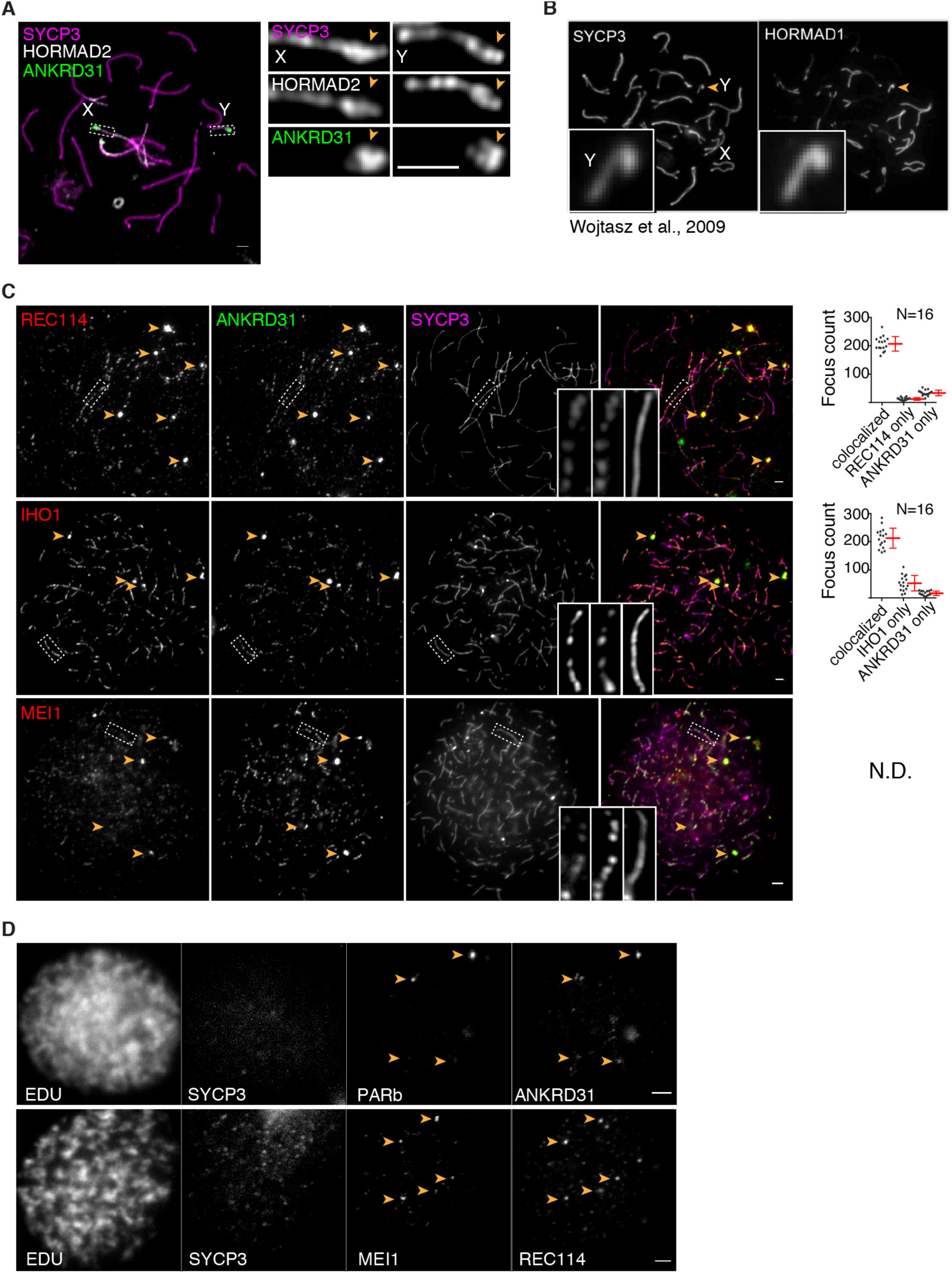
PAR axis thickening and accumulation of RMMAI proteins. **(A)** Axis thickening (SYCP3 and HORMAD2 staining) on the PAR (arrowhead) in a late zygotene spermatocyte. Scale bar: 2 μm. **(B)** Image adapted under Creative Commons CC-BY license from (*18*) showing enrichment of HORMAD1 on the thick PAR axis of the Y chromosome. **(C)** Colocalization of ANKRD31 and REC114, IHO1, and MEI1. Representative zygotene spermatocytes are shown. Arrowheads indicate densely staining blobs. Areas indicated by dashed boxes are shown at higher magnification at the right. The graphs show the total number of foci colocalized in leptotene/zygotene spermatocytes. N.D., not determined: The low immunofluorescence signal for MEI1 did not allow us to quantify the colocalization with ANKRD31, although MEI1 showed clear colocalization with ANKRD31 in the blobs and at least some autosomal foci (insets). Scale bars: 2 μm. Further evidence for extensive colocalization with ANKRD31 is documented in separate studies (*30, 31*)**. (D)** ANKRD31, REC114, and MEI1 immunostaining starts to appear in pre-leptonema. Seminiferous tubules were cultured with 5-ethynyl-3′-deoxyuridine (EdU) to label replicating cells, then chromosome spreads were stained for SYCP3 and either MEI1 plus REC114 or ANKRD31 plus PARb FISH. Colocalized foci appear in pre-leptonema (EdU-positive cells that are weakly SYCP3-positive), as previously shown for MEI4 and IHO1 (*21, 23*). Because we can already detect ANKRD31 accumulation at sites of PARb-hybridization, we infer that the stronger sites of accumulation of MEI1 and REC114 also include PARs. Scale bars: 2 μm.

**Fig. S2.**
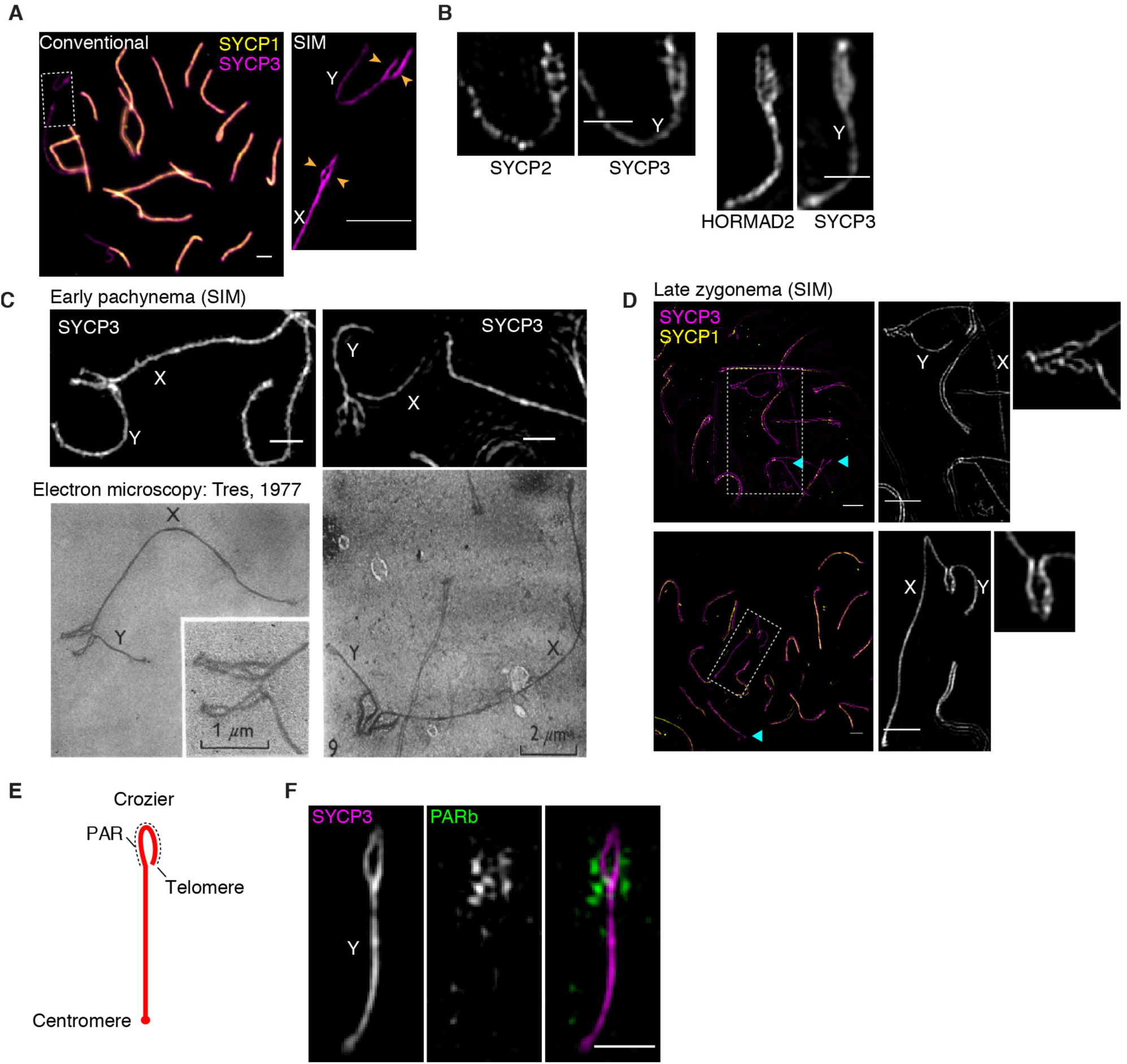
PAR ultrastructure. **(A)** Comparison of conventional microscopy and SIM, showing that the thickened PAR axis in conventional microscopy is resolved as separated axial cores (arrowheads). Scale bars: 2 μm. **(B)** Ultrastructure of axis proteins SYCP2, SYCP3, and HORMAD2 in the PAR. Scale bars: 1 µm. **(C,D)** Paired PARs with elongated and split axes occur in late zygonema to early pachynema. Shown are electron micrographs adapted with permission from (*34*) in comparison with SIM immunofluorescence images of spermatocytes at early pachynema (panel C; same chromosomes as in **Fig. 1E**) or late zygonema (panel D; cyan arrowheads indicate examples of incomplete autosomal synapsis). The spermatocytes in the electron micrographs were originally considered to be in mid-to-late pachynema (*34*). However, in our SIM experiments, we can only detect this structure (paired X and Y with elongated and split axes, resembling a crocodile’s jaws) around the zygotene–pachytene transition, when RMMAI proteins are still highly abundant on the PAR axes, and when most or all autosomes are completely synapsed. Moreover, other published electron micrographs from mid-to-late pachytene spermatocytes show diagnostic ultrastructural features that are not present in the electron micrographs reproduced here, including a short PAR axis length, multi-stranded stretches of axis on non-PAR portions of the X and Y chromosomes with excrescence of axial elements, and a clear thickening of autosomal telomeres (*28, 57*). These observations allow us to conclude definitively that the elongation and splitting of PAR axes are a hallmark of cells from late zygonema into early pachynema. Scale bars in SIM images: 1 μm in panel C, 2 µm in panel D. **(E)** Cartoon of hypothetical crozier configuration in which a single conjoined axis for both sister chromatids is folded back on itself. **(F)** Separation of PAR sister chromatid axes. FISH signal for the PARb probe is arrayed relatively symmetrically on both axial cores, consistent with separated sister chromatid axes (a bubble configuration). Scale bar: 1 µm.

**Fig. S3.**
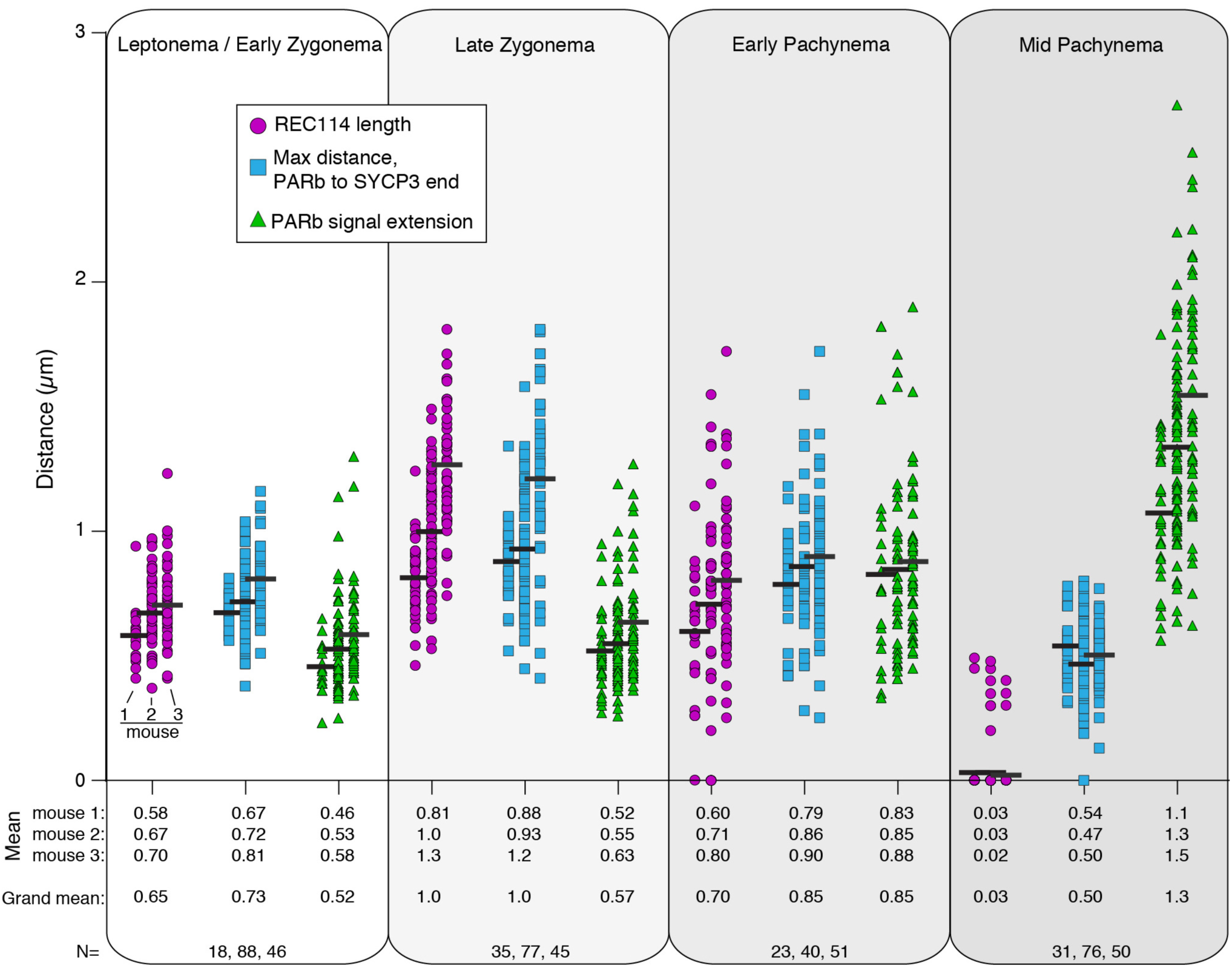
Time course of the spatial organization of the PAR loop–axis ensemble. Measurements of REC114 spreading along the PAR axis, length of the PAR axis, and extension of PARb chromatin orthogonal to the axis were collected as in **Fig. 2B**, on two additional males. Data from mouse 2 are reproduced from **Fig. 2B** to facilitate comparison. Means of each measurement for each mouse at each stage are given below, along with the means across all three mice. Means are rounded to two significant figures; the grand mean was calculated using unrounded values from individual mice. The number of cells of each stage from each mouse is given (N). Modest variability in the apparent dimensions of the Y chromosome PAR between different mice may be attributable to variation in copy number of mo-2 and other repeats because of unequal exchange during meiosis. Nonetheless, highly similar changes in spatial organization over time in prophase were observed in all mice examined, namely progressive elongation then shortening of axes and concomitant lengthening of loops.

**Fig. S4.**
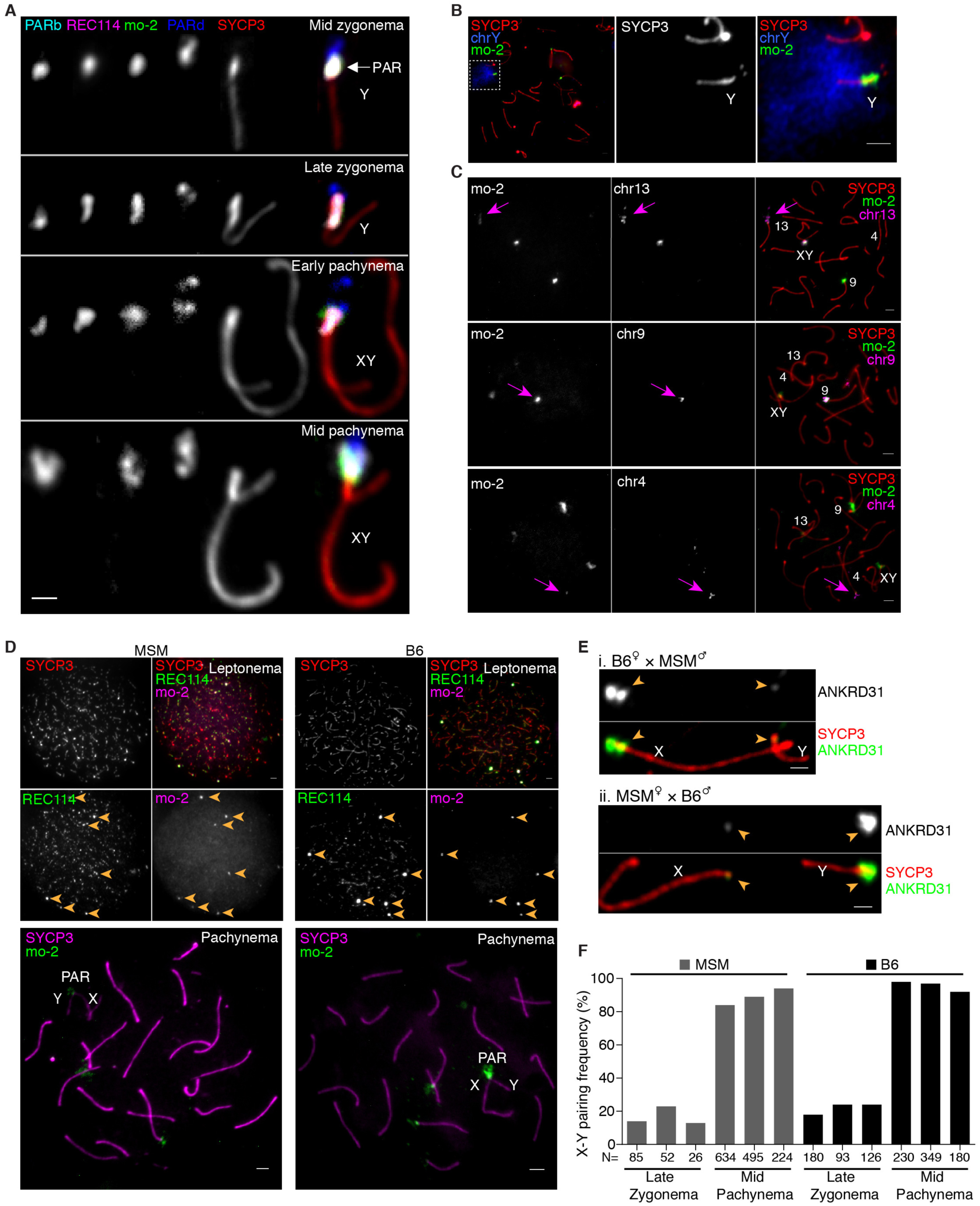
RMMAI enrichment at mo-2 minisatellite arrays in the PAR and on specific autosomes. **(A)** Comparison of mo-2 FISH with REC114 localization relative to the PAR boundary (PARb FISH probe) and the distal PAR (PARd probe). In mid zygonema, the mo-2 FISH signal colocalizes well with REC114 staining in between the PARb and PARd FISH signals. In late zygonema, mo-2 and REC114 are similar to one another and are elongated along the thickened SYCP3 staining of the PAR axis. From early to mid pachynema, REC114 progressively disappears, whereas the mo-2 FISH signal becomes largely extended away from the PAR axes. Note that the relative positions of the PARb and PARd probes reinforce the conclusion that the PAR does not adopt a crozier configuration. Scale bar: 1 μm. **(B)** Illustration of the compact organization of the PAR chromatin (mo-2 FISH signal) compared to a whole-Y-chromosome paint probe. Scale bar: 2μm. **(C)** Confirmation that autosomal mo-2 FISH signals match the chromosomal locations indicated by mm10 or Celera genome assemblies. FISH was performed using an oligonucleotide probe containing the mo-2 consensus sequence in combination with BAC probes for adjacent segments of chromosomes 13, 9 and 4, as indicated. Magenta arrows point to concordant FISH signals. The chr13 BAC probe also hybridizes to the PAR. Scale bars: 2 µm. **(D)** Lower mo-2 copy number in the *M. m. molossinus* subspecies correlates with lower REC114 staining in mo-2 regions. The top panels compare MSM and B6 mice for the colocalization between REC114 immunostaining and mo-2 FISH in leptotene spermatocytes. The REC114 and SYCP3 channels are shown at equivalent exposure for the two strains, whereas a longer exposure is shown for the mo-2 FISH signal in the MSM spermatocyte. Note that the mo-2-associated REC114 blobs are much brighter relative to the smaller dispersed REC114 foci in the B6 spermatocyte than in MSM. The bottom panel shows representative pachytene spermatocytes to confirm the locations of mo-2 clusters at autosome ends and the PAR in the MSM background. Scale bars: 2 μm. **(E)** Representative micrographs of late zygotene spermatocytes from reciprocal F1 hybrid males from crosses of B6 (high mo-2 copy number) and MSM (low mo-2 copy number) parents. Scale bar: 1 µm. **(F)** Frequency of paired X and Y at late zygonema and mid pachynema analyzed in three MSM and three B6 males. Differences between strains were not statistically significant at either stage (p > 0.13, Student’s t test). Note also that MSM X and Y are late pairing chromosomes, as in the B6 background. The similar pairing kinetics indicates that the lower intensity of RMMAI staining on the MSM PAR is not attributable to earlier PAR pairing and synapsis in this strain. The number of spermatocytes analyzed is indicated (N).

**Fig. S5.**
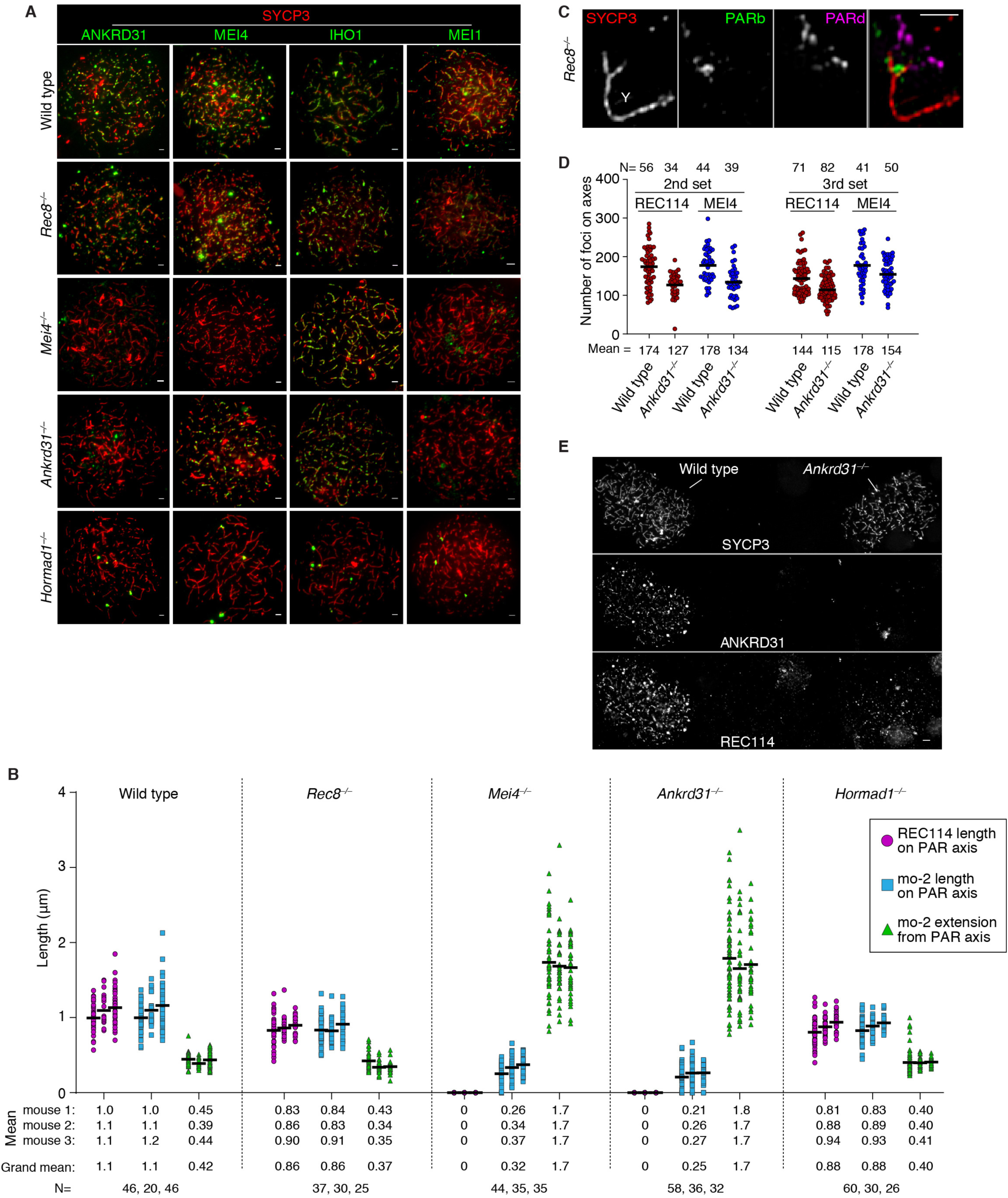
Genetic requirements for RMMAI assembly on chromosomes and for PAR loop–axis organization. **(A)** Representative micrographs of ANKRD31, MEI4, IHO1 and MEI1 staining in wild type and the indicated mutants (quantification is in **Fig. 4A**). Scale bars: 2 μm. **(B)** Measurements of PAR loop–axis organization, as in **Fig. 4B**, on two additional males. Data from mouse 1 are reproduced from **Fig. 4B** to facilitate comparison. Means of each measurement for each mouse at each stage are given below, along with the means across all three mice. Means are rounded to two significant figures; the grand mean was calculated using unrounded values from individual mice. The number of cells of each stage from each mouse is given (N). **(C)** REC8 is dispensable for splitting apart of PAR sister chromatid axes, but is required to maintain the connection between sisters at the distal tip of the chromosome. A representative SIM image is shown of a Y chromosome from a late zygotene *Rec8*^*–/–*^ mutant spermatocyte. The SYCP3-labeled axes adopt an open fork configuration. Note that the distal FISH probe (PARd) shows that there are clearly disjoined sisters whereas the PAR boundary (PARb) shows only a single compact signal comparable to wild type. The disposition of the probes and SYCP3 further rules out the crozier configuration as an explanation for split PAR axes. Scale bar: 1 µm. **(D)** Quantification of REC114 and MEI4 foci in two additional pairs of wild type and *Ankrd31*^*–/–*^ mice. Horizontal lines indicate means. Fewer foci were observed in the *Ankrd31*^*–/–*^ mutant (p < 0.02, Student’s t test) for each comparison of mutant to wild type. **(E)** Reduced REC114-staining intensity of axis-associated foci in *Ankrd31*^*–/–*^ mutants. To rigorously control for slide-to-slide and within-slide variation in immunostaining, we mixed together wild-type and *Ankrd31*^*–/–*^ testis cell suspensions before preparing chromosome spreads. A representative image is shown of a region from a single microscopic field containing two wild-type zygotene spermatocytes (left) and two *Ankrd31*^*–/–*^ spermatocytes of equivalent stage (right). Note the diminished intensity of REC114 foci in the *Ankrd31*^*–/–*^ spermatocytes. Scale bar: 2 µm.

**Fig. S6.**
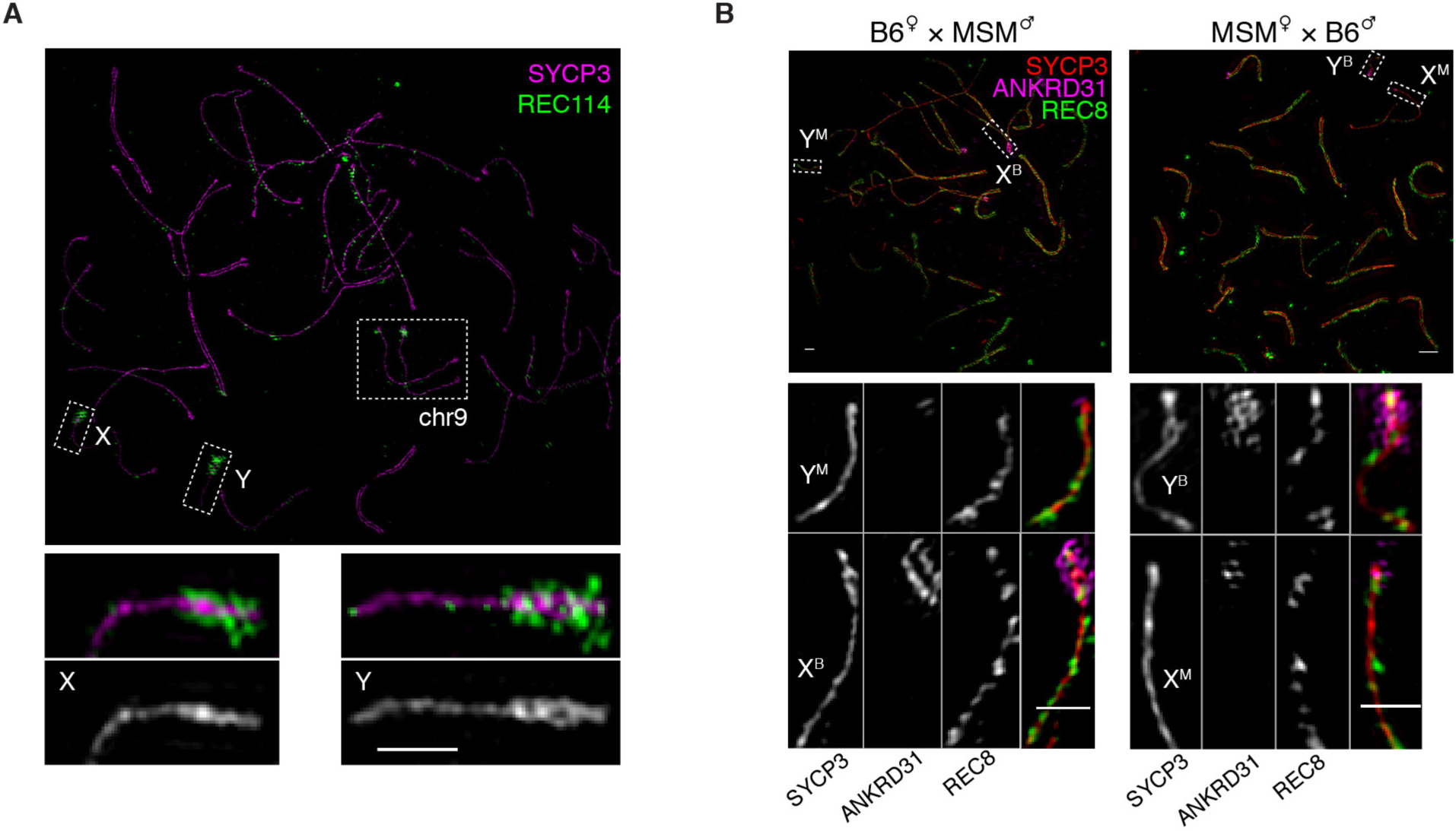
RMMAI recruitment and axis remodeling are functionally connected to mo-2 presence and copy number. **(A)** The mo-2-containing portion of chr9 undergoes axis elongation and splitting similar to PARs. The upper panel shows the SIM image for the complete wild-type zygotene spermatocyte from which the higher magnification detail of chr9 (identified based on chromosome length and brightness of REC114 staining) is shown in **Fig. 5A**. Higher magnification views of the Y and PAR-containing part of the X are shown for comparison. Scale bar: 1 µm. **(B)** Low mo-2 copy number on *M. m. molossinus* sex chromosomes is associated with a lesser degree of loop–axis reorganization. Representative SIM images are shown of late zygotene spermatocytes from F1 hybrid males from reciprocal crosses of B6 (high mo-2 copy number) and MSM (low mo-2 copy number) parents. Scale bars: 1 µm.

**Fig. S7.**
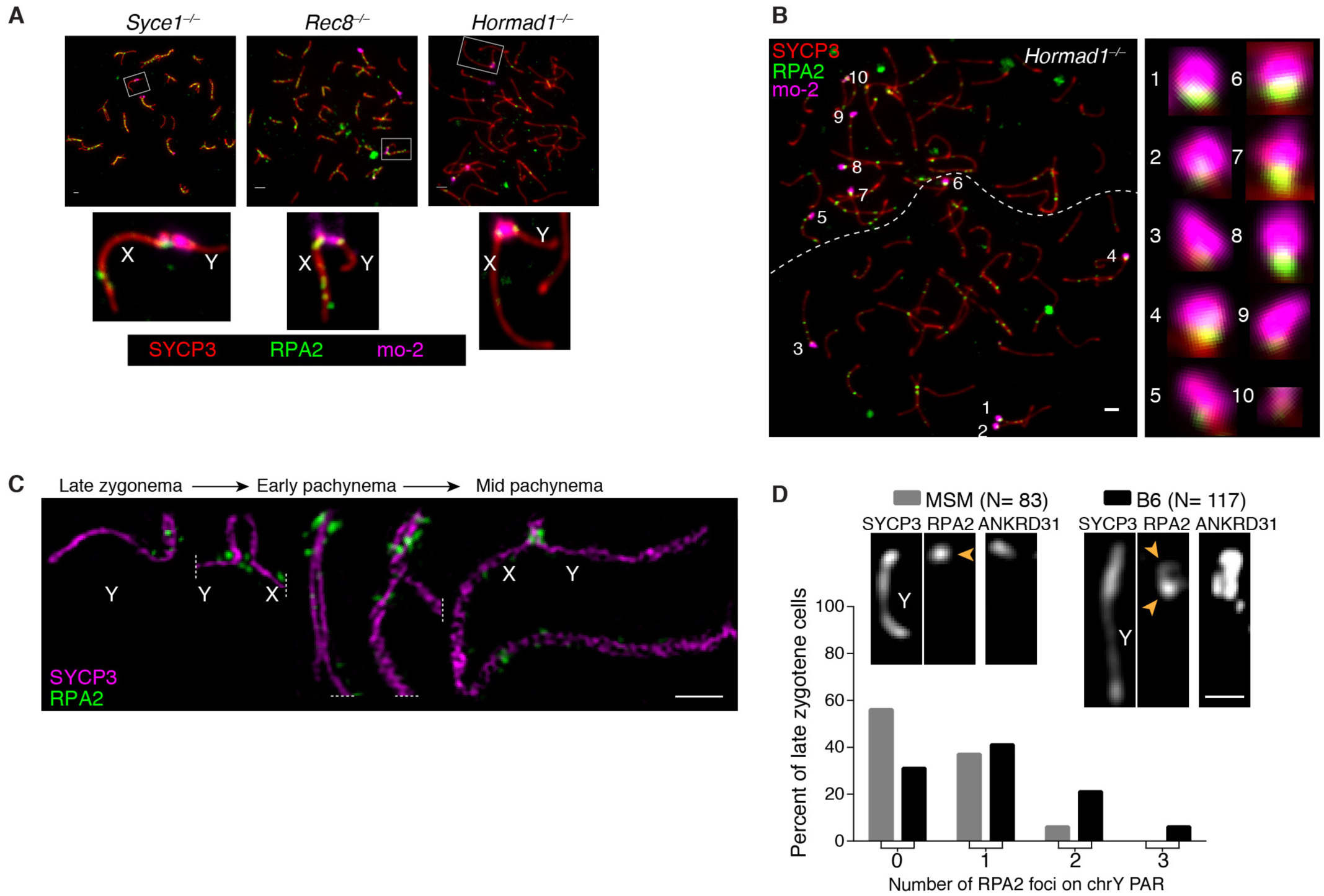
PAR-associated RPA2 foci. **(A)** Immuno-FISH for RPA2 and mo-2 was used to detect DSBs cytologically in wild type and the indicated mutants. Representative images are shown for additional mutants, supplementing the images in **Fig. 5C**. Quantification is in **Fig. 5D,E**. Scale bars: 2 µm. **(B)** Frequent DSB formation at mo-2 regions in the PARs and on autosomes does not require HORMAD1. Micrograph at left shows two adjacent spermatocytes (boundary indicated by dashed line). Insets at right show higher magnification views of the numbered mo-2 regions, all of which are associated with RPA2 immunostaining of varying intensity. Scale bar: 2 µm. **(C)** Montage of SIM images from a B6 male showing that multiple, distinct RPA2 foci can be detected from late zygonema to mid pachynema, suggesting that multiple PAR DSBs can be formed during one meiosis (see also (*7*) for further discussion). Scale bar: 1 μm. **(D)** Percentage of spermatocytes at the zygotene-pachytene transition with no (0), 1, 2 or 3 distinguishable RPA2 foci on the unsynapsed Y chromosome PAR of MSM and B6 mice. The difference between the strains is statistically significant (negative binomial regression, p = 7.2 × 10^−5^). N indicates the number of spermatocytes analyzed. A representative picture is shown for each genotype, with one RPA2 focus on the MSM PAR and two apparent sites of RPA2 accumulation on the B6 PAR. Scale bar: 1 μm.

**Fig. S8.**
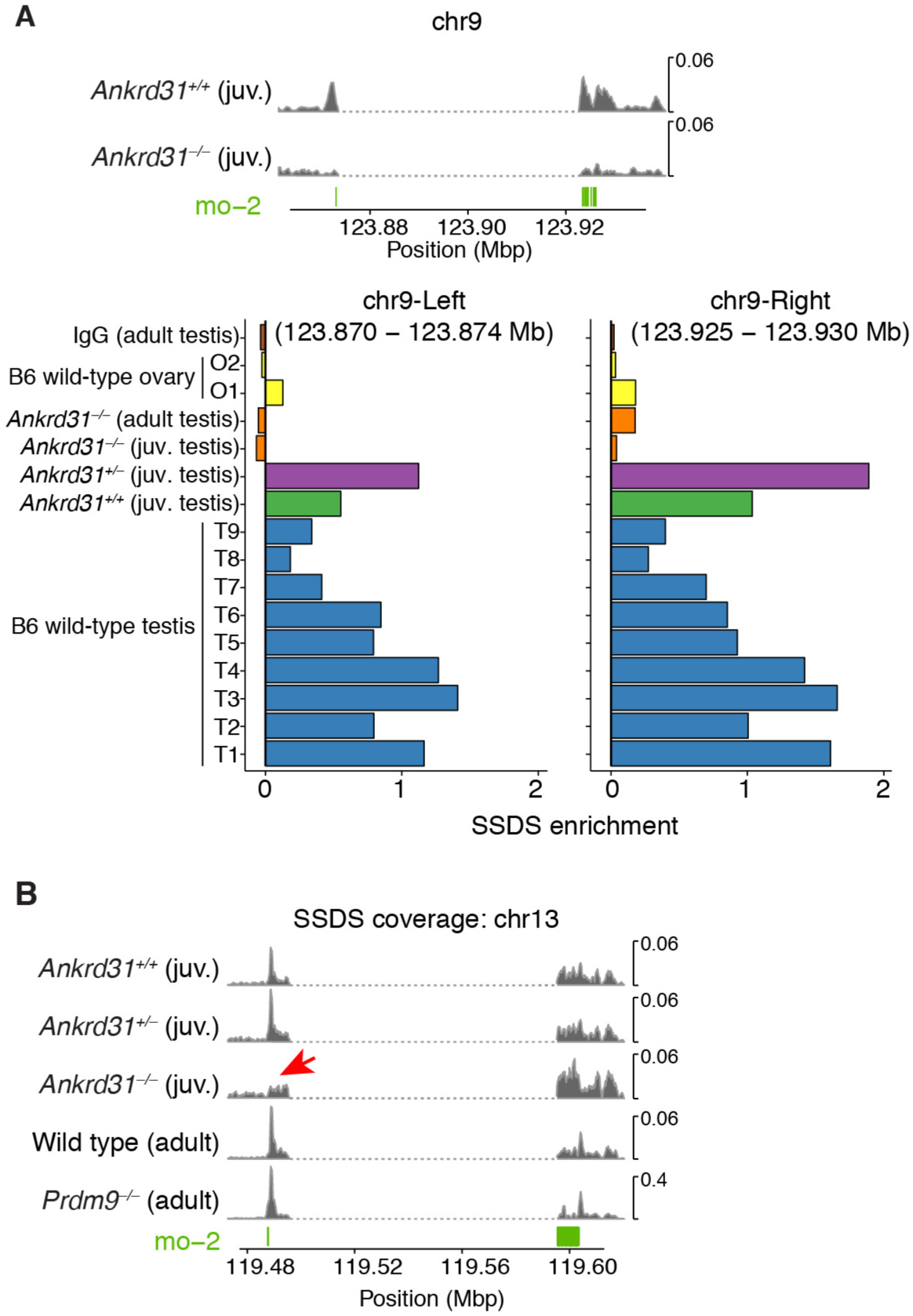
DSB maps on the PAR and autosomal mo-2 regions. **(A)** Regions adjacent to the mo-2 region on chr9 show SSDS signal that is reproducibly elevated relative to chr9 average in wild-type testis samples but not in maps from *Ankrd31*^*–/–*^ testes or wild-type ovaries. Two of the SSDS browser tracks are reproduced from **Fig. 5G**. Enrichment values are defined as coverage across the indicated coordinates relative to mean coverage for chr9 (see Methods for details). Note that ovary sample O1 and the *Ankrd31*^*–/–*^ adult sample are known to have poorer signal:noise ratios than the other samples (*30, 43*). **(B)** SSDS maps for the mo-2 region on chr13. The red arrow indicates an ANKRD31-dependent, PRDM9-independent peak. The dashed segment indicates a gap in the mm10 genome assembly. ***Note:*** For all SSDS coverage tracks in this study, reads mapping to multiple locations are included after random assignment to one of their mapped positions. However, the same conclusions are reached about ANKRD31-dependence and PRDM9-independence of signal on chr9 and chr13 if only uniquely mapped reads are used (data not shown).

**Fig. S9.**
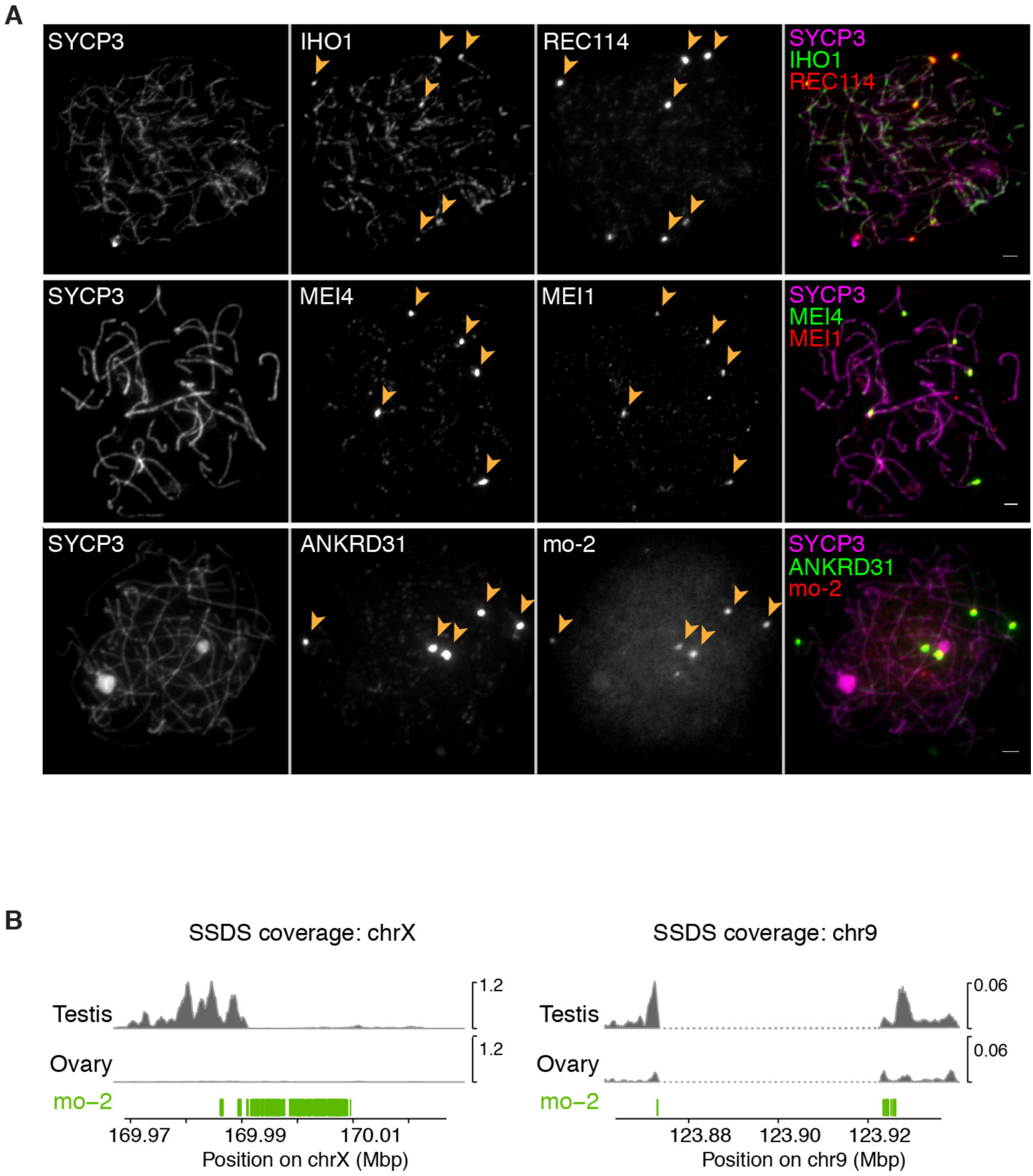
RMMAI accumulation and high-level DSB formation on mo-2 regions in oocytes. **(A)** Examples of zygotene oocytes showing the colocalization between blobs of IHO1 and REC114, MEI4 and MEI1, ANKRD31 and mo-2 FISH signal. Scale bars = 2 µm. **(B)** Substantially less DSB formation occurs in oocytes near the mo-2 region on chr9. SSDS signal is from (*43*). The X-PAR is shown for comparison [previously shown to be essentially devoid of DSBs in ovary samples (*43*)]. See **fig. S8A** for quantification.

**Fig. S10.**
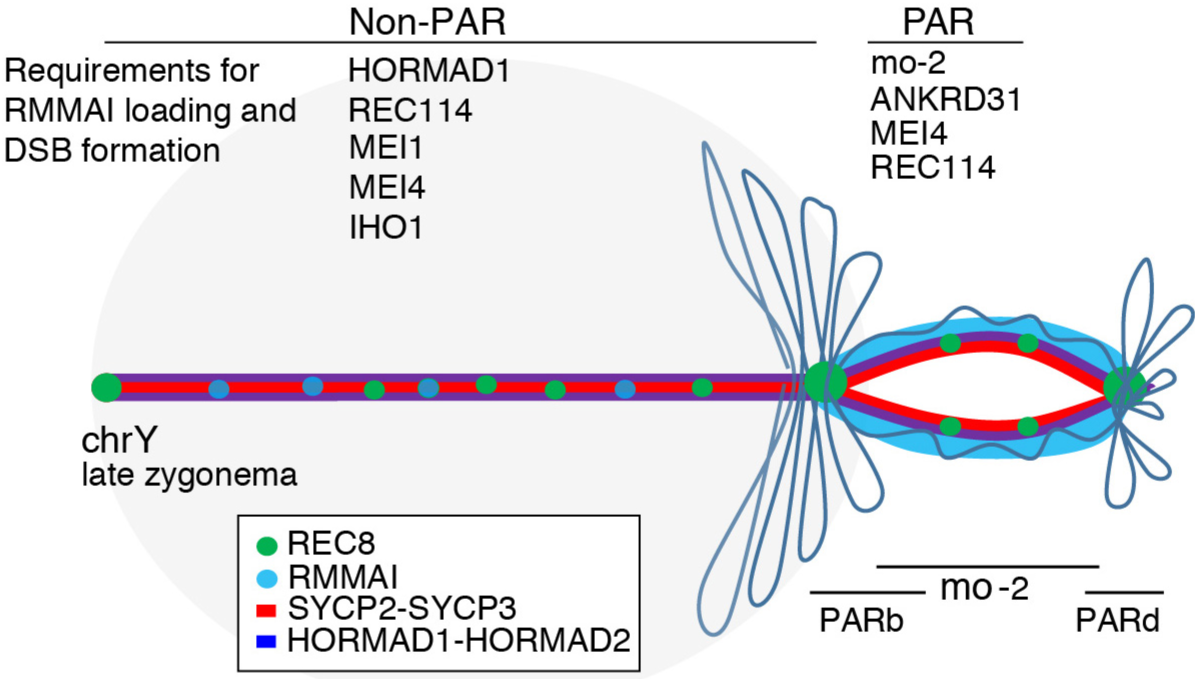
Summary of PAR ultrastructure and molecular determinants of axis remodeling and DSB formation. Schematic representation of the meiotic Y chromosome loop/axis strucuture before X–Y pairing at the transition between zygonema and pachynema. The chromosome axis comprises the meiosis-specific axial proteins SYCP2, SYCP3, HORMAD1, and HORMAD2; cohesin subunits (only REC8 is represented); and the RMMAI proteins. On the non-PAR portion of the Y chromosome axis (left), RMMAI protein loading and DSB formation are partly dependent on HORMAD1 and ANKRD31, and strictly dependent on MEI4, REC114 (*24*), IHO1 (*31*), and presumably MEI1 (*29*). The DNA is organized into large loops, with a low number of axis-associated RMMAI foci. By contrast, in the PAR (right), the hyper-accumulation of RMMAI proteins at mo-2 minisatellites (possibly spreading into adjacent chromatin) promotes the elongation and subsequent splitting of the PAR sister chromatid axes. Short mo-2-containing chromatin loops stretch along this extended PAR axis, increasing the physical distance between the PAR boundary and the distal PAR sequences, including the telomere. The degree of RMMAI protein loading, PAR axis differentiation, and DSB formation are proportional to the mo-2 FISH signal (which we interpret as reflecting mo-2 copy number), and depend on MEI4 and more specifically ANKRD31.

